# Random walk and cell morphology dynamics in *Naegleria gruberi*

**DOI:** 10.1101/2023.08.25.554756

**Authors:** Masahito Uwamichi, Yusuke Miura, Ayako Kamiya, Daisuke Imoto, Satoshi Sawai

## Abstract

Amoeboid cell movement and migration are wide-spread across various cell types and species. Microscopy-based analysis of the model systems *Dictyostelium* and neutrophils over the years have uncovered generality in their overall cell movement pattern. Under no directional cues, the centroid movement can be quantitatively characterized by their persistence to move in a straight line and the frequency of re-orientation. Mathematically, the cells essentially behave as a persistent random walker with memory of two characteristic time-scale. Such quantitative characterization is important from a cellular-level ethology point of view as it has direct connotation to their exploratory and foraging strategies. Interestingly, outside the amoebozoa and metazoa, there are largely uncharacterized species in the excavate taxon Heterolobosea including amoeboflagellate *Naegleria*. While classical works have shown that these cells indeed show typical amoeboid locomotion on an attached surface, their quantitative features are so far unexplored. Here, we analyzed the cell movement of *Naegleria gruberi* by employing long-time phase contrast imaging to automatically track individual cells. We show that the cells move as a persistent random walker with two time-scales that are close to those known in *Dictyostelium* and neutrophils. Similarities were also found in the shape dynamics which is characterized by the appearance, splitting and annihilation of the curvature waves along the cell edge. Our analysis based on the Fourier descriptor and a neural network classifier point to importance of morphology features unique to *Naegleria* including complex protrusions and the transient bipolar dumbbell morphologies.

## INTRODUCTION

Combinatorial use of persistent motion and reorientation is a common feature found in cell movement. Be it bacterial swimming or amoeboid crawling, persistent movement allows cells to gain most distance in one preferred direction so as to facilitate efficient escape from hazards or conversely attraction to nutrients. Reorientation on the other hand is not only required to adjust direction of persistent movement but also to facilitate cells to randomly explore and survey uncertain extracellular environments (Viswanathan et al., 1999; Bartumeus et al., 2002). In *Eschrichia coli* bacteria, the cell movement consists of a period of straight run interrupted by a stall or “tumble” where flagellar rotation reverses and cells reorient in random directions. The frequency of tumbling is regulated through a chemosensory system so as to provide orientation bias towards an attractant or away from a repellent. The exact nature of such motility pattern determines how well *E. coli* cells disperse (Taktikos et al., 2013). In the amoeboid movement, pseudopodal protrusions enriched in branched F-actin networks (Pollard, 2007) are formed transiently and can guide cells in different orientations. In addition, a confined region of the plasma membrane needs to retract in order to realize net displacement. In many cell types, cortical F-actin that is crosslinked with myosin II are enriched in such contractile membrane regions (Chi et al., 2014). Persistent movement arises when a cell has mono-polarity meaning that it has a single dominating leading edge and a retracting trailing end. When and where along the plasma membrane these organizations occur determine the ordered occurrence of protruding plasma membrane and rear retraction thus ultimately dictate the direction, speed and duration of cell movements.

Quantitative time-series analyses of cell displacement and cell shape change are important for explicit characterization of random cell motion. In many cases, cell displacement can be approximated as a particle obeying persistent random walk. Phenomenologically, the simplest form of differential equation that describes such stochastic dynamics is the Langevin equation (Dunn and Brown, 1987; Selmeczi et al., 2005, 2008).

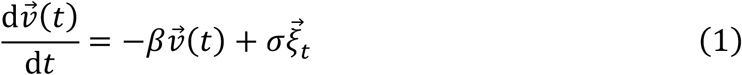

Where 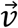 is the velocity vector, *β* is the memory coefficient,*σ* is the noise strength, and 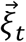 is 2D white Gaussian noise. Random walk of *E. coli* can be approximated by Brownian motion having a short-term memory. In eukaryotic crawling, cell trajectories of fibroblast cells (Gail and Boone, 1970) and endothelial cells (Stokes et al., 1991) are also consistent with this simple persistent random walk model. In many other cell types, random walk includes memory that depends on the velocity and orientation which can be described by modifications to the above model (Takagi et al., 2008; Li et al., 2011). There are also random walk statistics called Levy-walk with step lengths that follows a long tail (power-law) distribution (Viswanathan et al., 1999). There, the Mean Square Displacement (MSD) essentially diverges and the trajectories are characterized by self-similarity of the step lengths (Reynolds, 2010; Reynolds and Ouellette, 2016). Because Levy-walk has very small probability of revisiting the same location, it is thought to arise in systems such as in bird foraging that require an efficient search strategy. At the cellular-level, effector T-cells (Harris et al., 2012), swarming bacteria have been reported to exhibit Levy-flight like statistics (Ariel et al., 2015; Huo et al., 2021).

To date, quantitative understanding of random walk behavior of amoeboid cells is limited to data from a handful of cell-types; these are mostly timelapse microscopy images of cultured metazoan cells and amoebozoa *Dictyostelium*. From microbial ethology and evolutionary biology perspectives, however, we should note that amoeboid movement is found not only in animals, fungi and amoebozoa (Prostak et al., 2021) but also in largely uncharacterized species in the excavate taxon Heterolobosea namely Naegleria spp. and the slime mold acrasids (Brown et al., 2012). The ancestors of the opisthokont lineage and *Naegleria* diverged more than a billion years ago (Parfrey et al., 2011). Knowing the details of motility characteristics in *Naegleria* should help us understand the common design of the motility trait that is either deeply conserved across taxa or acquired independently by strong selective advantages. Additionally, there are biomedical implications in studying *Naegleria* cell motility. Naegleria species including *Naegleria fowleri* are one of several known ‘brain-eating’ amoebae that cause fatal central nervous system infection called amebic meningoencephalitis. Their pathogenicity is thought to be related to their capacity to enter brain by penetrating nasopharygeal mucosa and migrate along olfactory nerves (Thong and Ferrante, 1986).

Among members of genus *Naegleria*, non-pathogenic *Naegleria gruberi* is the better characterized species whose genome has been sequenced (Fritz-Laylin et al., 2010). In its amoebic phase, *Naegleria gruberi* grows and divides by feeding on bacteria through phagocytosis (Fulton, 1970). Under low electrolyte conditions, it quickly shifts to the non-feeding flagellated state by rapid de novo synthesis of microtubles (Walsh, 2007). In the amoebic state, the overall cell cortex is enriched in F-actin with marked accumulation around membrane ruffles (Velle and Fritz-Laylin, 2020). An early work using reflection interference microscopy have revealed that *Naegleria gruberi* adhere and form discrete dot-like contacts to non-treated glass surfaces and migrate (Preston and King, 1978). These so-called ‘focal contacts’ leave behind footprints of membrane residues on the glass substrate as the cells crawl away (Preston and King, 1978). With the advent of genomics and molecular cell biology, it has become clear that *Naegleria gruberi* possess the essential side-branching nucleator of F-actin - the Arp2/3 complex and its activators WASP and SCAR (Fritz-Laylin et al., 2017; Velle and Fritz-Laylin, 2020; Prostak et al., 2021). Inhibition of Formin reduces directional persistence, and inhibition of the Arp2/3 complex reduces the cell speed (Velle and Fritz-Laylin, 2020). *Naegleria gruberi* also has Myosin II (Sebé-Pedrós et al., 2014) and a potential orthologue of Integrin beta (Morales et al., 2022), although whether they exist in other groups in excavata is unclear (Velle and Fritz-Laylin, 2019). While these works indicate likely similarity of actin-dependent processes involved in cell crawling in an evolutionary distant eukaryote, quantitative characterization of the cell-level motility pattern is so far lacking. Do *N. gruber*i cells exhibit persistent random walk behavior? What are the characteristic time scale of persistence and reorientation if any? How similar are their movements compared to the well-studied systems such as *Dictyostelium* and immune cells? In this report, we performed quantitative analysis on cell movements and shape change of *Naegleria gruberi*. Our analysis demonstrate that *Naegleria* cells exhibit persistent random walk driven by a large morphology change that involves appearance, splitting and annihilation of uniquely complex pseudopodium protrusions.

## RESULTS

To quantitate the movement of *Naegleria gruberi* on a two-dimensional flat surface, we performed phase contrast time-lapse microscopy. A non-coated glass coverslip was employed as a cell substrate throughout this study. **Figure 1A** shows representative phase contrast images of *Naegleria gruberi* in liquid growth media (Materials and Methods). The cells under our culture condition exhibited one or more hyaline protrusions that appeared dark in phase contrast images (**Figure 1A** arrows). In the example shown, protrusions extended along the glass surface for 15-50 seconds and the one that became dominant (i.e. the leading edge) extended in the direction of the overall cell movement (**Figure 1A**, 0 sec). Marked cytoplasmic streaming from the center of the cell towards these extensions was observed (Supplementary Movie S1). A new protrusion appeared and extended first in the lateral direction (**Figure 1A** 20 sec, 60sec arrow) and steered towards the front then was bent sideway until it was retracted (**Figure 1A** 40sec, 80 sec). Duration of the pseudopod extension/retraction cycle varied between 15 - 50 seconds (**Supplementary Figure S1; Supplementary Movie S2**). Concomitant reversal of cytoplasmic streaming was observed during retraction of pseudopods. A small bud-like bulge at the trailing end of a cell which we shall refer to as ‘uroid’ appeared as a residue of a retracted pseudopod that was retained for an extended period of time (**Figure 1A** white circle). The uroid contained thin filopodia-like projections as described earlier (Preston and King, 1978).

**FIGURE 1.**
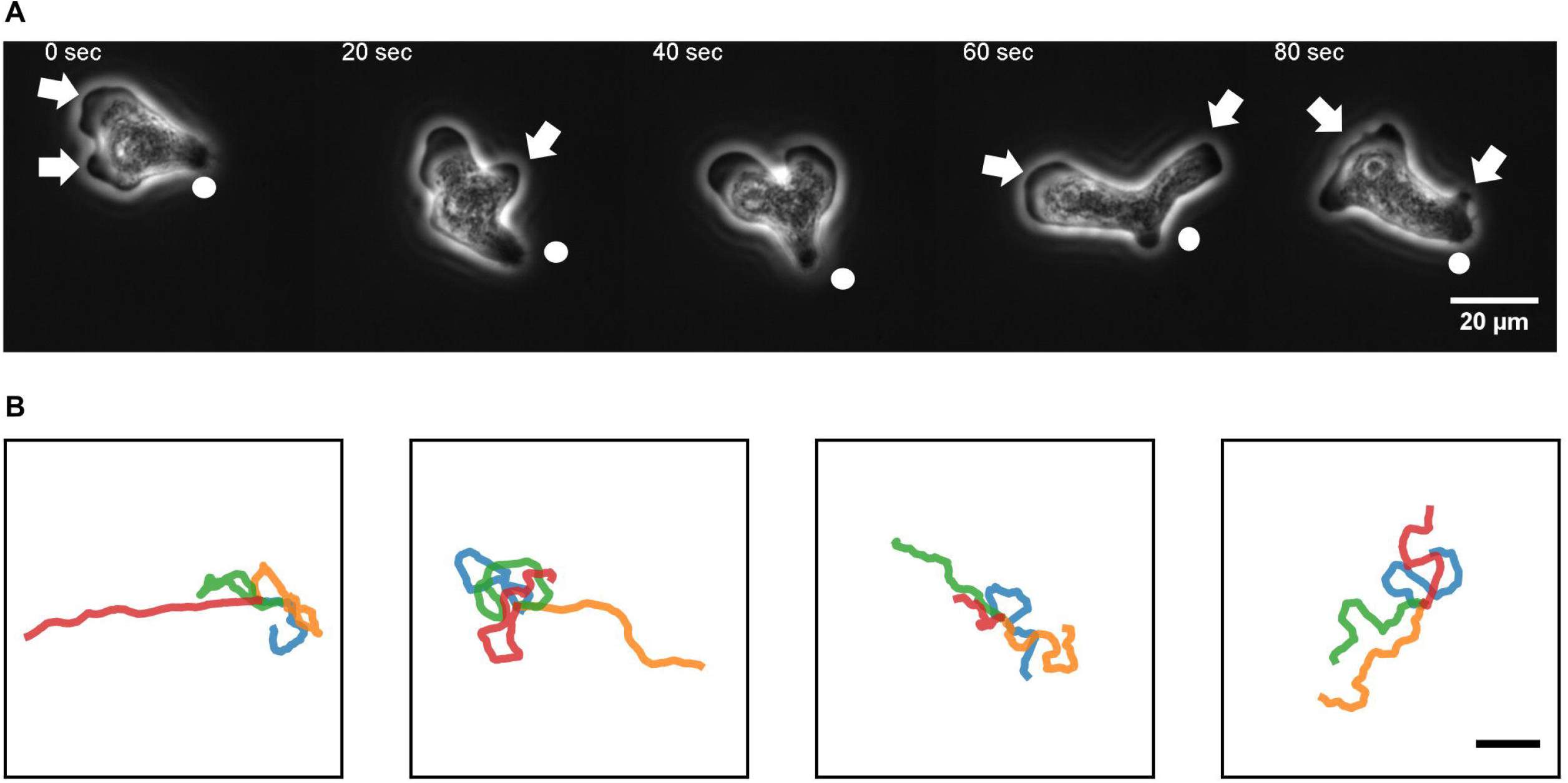
An overview of *N. gruberi* movement. **(A)** Representative phase contrast images from time-series of a migrating *N. gruberi* cell. Arrows: protruding edges. Circles: a bud-like rear structure (‘uropod’). **(B)** First 360 seconds of randomly selected centroid trajectories. 4 trajectories are separately shown for visibility. Scale bar: 100 μm.

Under our culture condition, the cells appeared to re-orient in random directions at irregular timing. We performed long time cell tracking by employing an automated stage that was programmed to track target cells (see Methods). **Figure 1B** shows representative cell trajectories obtained from the automated tracking. The trajectories consisted of a period of straight movement that lasted for about 30-200 seconds and a time period of relative low displacement and re-orientation (**Figure 1B**). The movement is thus, at surface, akin to the run-and-tumble behavior of *E. coli*. There was a close link between the run and re-orientation dynamics with the cell shape. During the straight run, the cells took a fan-like shape (**Figure 2A**; **Supplementary Movie S3**). The tail remained narrow while at the front was a broad lamellipodia that expanded then split into branches of pseudopods (**Figure 2A**, 0-16 sec). These bifurcating protrusions often fused to restore a large lamellar extension (**Figure 2A**, 20 sec). On the other hand, cells re-oriented when the bifurcated protrusions remained separate. In most cases, the uropod persisted during front splitting and thus the cells took a Y-or trident shape (**Figure 2B**; **Supplementary Movie S4**). There were also cases where the uroid disappeared in Y-shaped cells (**Figure 2C**; **Supplementary Movie S5**). The two fronts expanded in the opposing directions and gave rise to a transient ‘dumbbell-like’ bipolar morphology (**Figure 2C**, 70 sec). After 10 seconds, one end shrunk and became the uroid while the other end became the next front (**Figure 2C**, 80 sec). There was little centroid displacement during this period which lasted for about 40 seconds.

**FIGURE 2.**
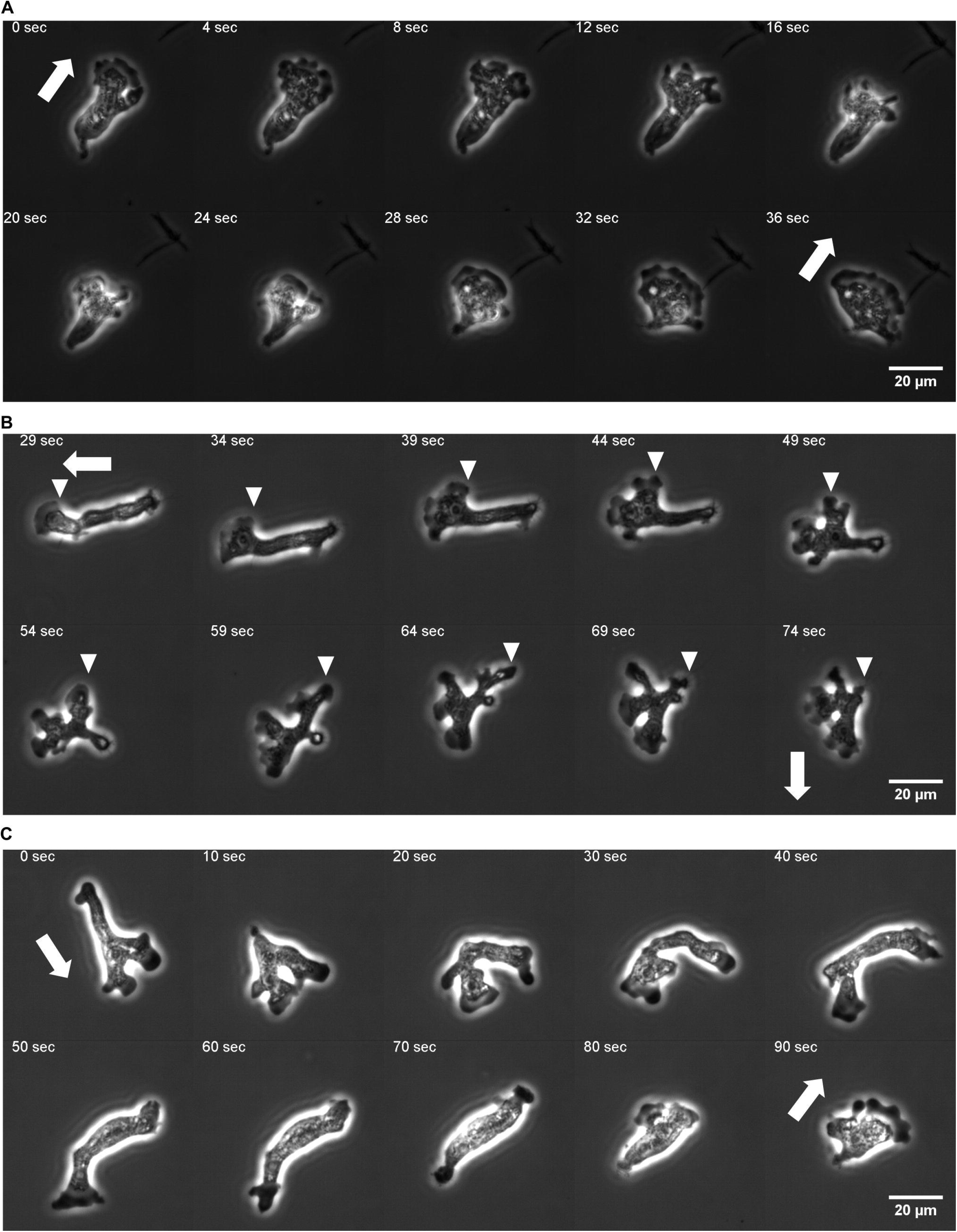
Protrusion dynamics and the cell shape change. **(A)** A fan-shaped cell with front splitting during persistent run. (**B**) Front splitting followed by reorientation (curvature kymograph for the sequence is shown in **Figures 5A,B**). (**C**) Dumbbell shape arise after front splitting and disappearance of the uropod (curvature kymograph for the sequence is shown in **Supplementary Figures S4D**,**F**). Arrows: orientation of centroid movement. Inverted triangles: propagating curvature waves.

To characterize the random-walk statistics, we quantified the <u>m</u>ean <u>s</u>quare <u>d</u>isplacement (MSD) and the instantaneous speed defined by the distance of centroid displacement in 1 sec time interval from trajectories of N = 10 cells (**Figure 3**). Even with the help of automated stage tracking, fast movement of *N. gruberi* made it difficult to track cells for long time duration before they come close to the edge of the plate or collided with one another. Thus, to obtain MSD, single trajectories were each divided into sub-trajectories of 100-3600 sec time-window and treated as independent data samples (**Figure 3A**). The slope of the MSD from the individual trajectories was 1.5 to 2.0 (**Supplementary Figure S2A**). The slope of the ensemble-averaged MSD was 1.8 (**Figure 3B**). The time-dependency of the MSD indicates that the random walk of *Naegleria* falls somewhere between pure Brownian (exponent of 1) and ballistic (constant velocity) motion (exponent of 2). **Figure 3C** and **Supplementary Figures S2B-F** show the distribution of the instantaneous velocity. The distribution followed 2-dimensional Gaussian with zero-mean and standard deviation of 51 μm/min (**Figure 3C, Supplementary Figures S2B-F**). This feature is distinct from *Dictyostelium* random motility which is non-Gaussian (Takagi et al., 2008). The median of the absolute speed was 60 μm/min which is close to the average speed reported in earlier literatures (King et al., 1981; Thong and Ferrante, 1986).

**FIGURE 3.**
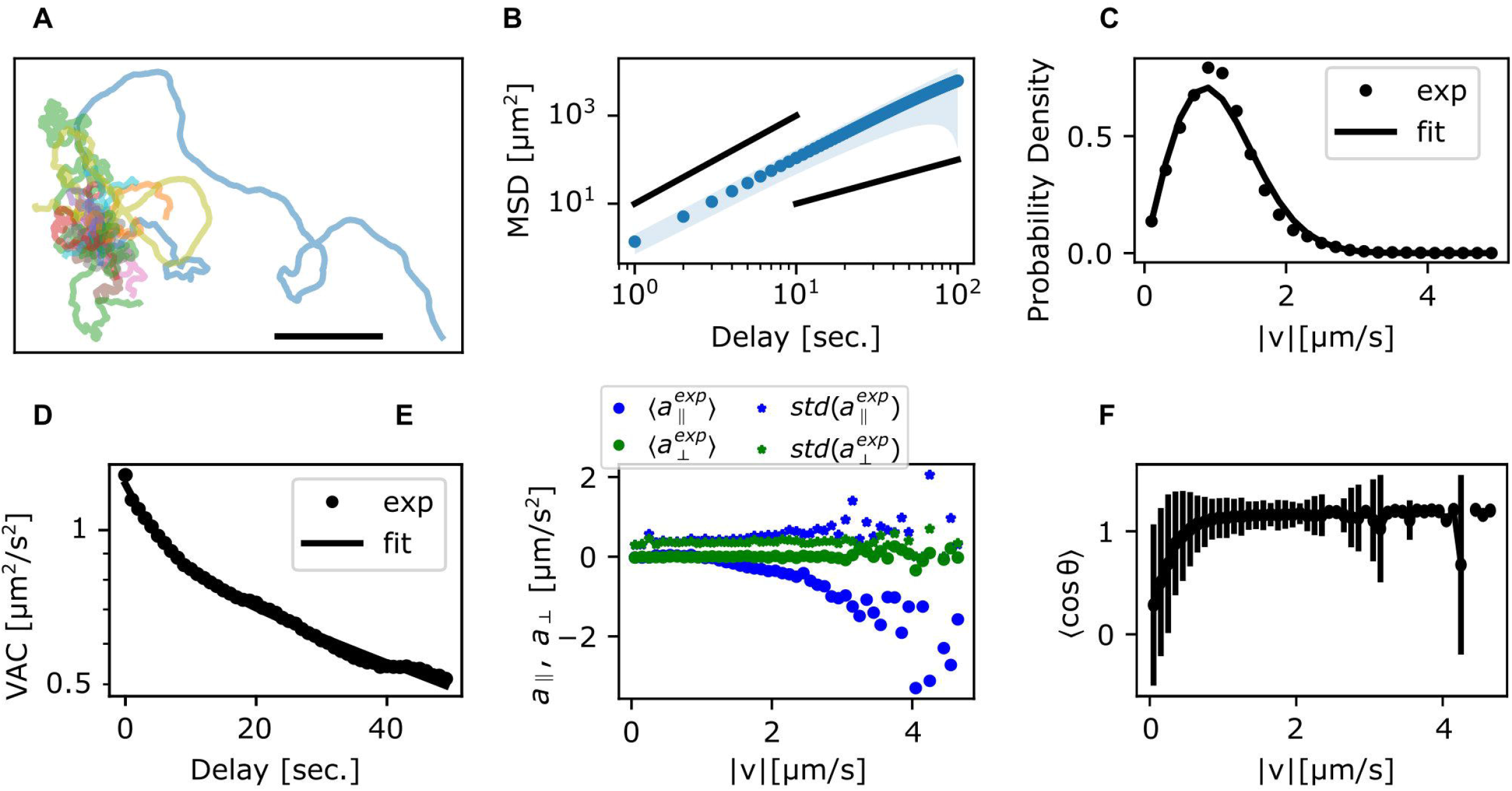
Statistics of the centroid movement. **(A)** The trajectories used for detailed analysis (N = 35). The origin is set at the initial position. Scale bar: 500 μm. **(B)** The ensemble averaged MSD. Solid black lines: exponent 1 and 2. Shaded region: standard deviation. **(C)** Probability distribution of 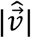. Solid black: a fitting curve from a 2-dimensional isotropic Gaussian distribution with a standard deviation of 0.86 μm/sec. **(D)** The ensemble averaged VAC. Black solid curve: the double exponential function (Eq. 2). **(E)** Acceleration parallel (blue) and orthogonal (green) to the velocity. The binned average (circle) and the standard deviation (star). **(F)** Persistence of displacement orientation in a unit time step. Cosine of the angle change θ in the velocity in 1 second interval. The binned average (circles) and the standard deviation (error bars).

The temporal auto-correlation of the centroid speed (<u>v</u>elocity <u>a</u>uto-<u>c</u>orrelation; VAC) (**Supplementary Figure S2G**) shows, on average, that there are two characteristic decay time that cross over at around 10 sec (**Figure 3D**). By assuming that VAC follows the sum of two exponential function (Selmeczi et al., 2005):

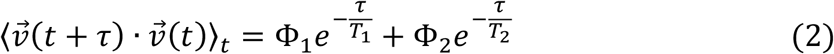

we obtained the decay time *T*_1_ and *T*_2_ of approximately 6 sec and 90 sec, respectively, where the weight Φ_1_ and Φ_2_ are 0.36 μm^2^/s^2^ and 0.87 μm^2^/s^2^ (**Figure 3D** black curve). To check for orientational preference in the memory, we plotted the relationship between velocity and acceleration (change in |*v*| in 1 sec interval) (**Supplementary Figure S2H**,**I**). The mean acceleration orthogonal (*a*_⊥_) to the velocity was near zero regardless of |*v*| (**Figure 3E**; green circle) with non-zero variance (**Figure 3E** green stars) which suggests that the orientation of *N. gruberi* has no apparent left-right asymmetry. On the other hand, the mean acceleration parallel (*a*_∥_) to the velocity was near zero at small velocity then decreased towards the negative at large velocities (**Figure 3E**). The standard deviation (**Figure 3E**; blue stars) increased somewhat at high |*v*|, however rarity of these fast step events prevented us from obtaining reliable averages. These features of acceleration are similar to those reported for *Dictyostelium* (Takagi et al., 2008). The negative acceleration parallel to the migration direction at high |*v*| implies that the cells do not maintain high |*v*| during re-orientation. Rather, movement in one direction is decelerated so that the non-memory term i.e. fluctuating components plays a dominant role in determining the next move. Accordingly, when we plot reorientation angle *θ* as a function of |*v*| (**Figure 3F**) we see that most of re-orientation occurs below |*v*| =1 μm/sec. Above 1 μm/sec the cells are moving in a straight line; i.e. cosθ = 1.

To gain further insights on the specifics of the random walk statistics, it is instructive to compare the data with the behavior of simple idealized equations. The velocity auto-correlation that follows the sum of two exponential indicates that random walk dynamics cannot be captured simply by the Ornstein-Uhlenbeck process (**Eq. 1**) which only has a single exponent (Dunn and Brown, 1987). A straight-forward and minimal extension to **Eq. 1** is to include additional memory with the decay rate *γ* as an integral in the form of generalized Langevin-equation (Selmeczi et al., 2005)

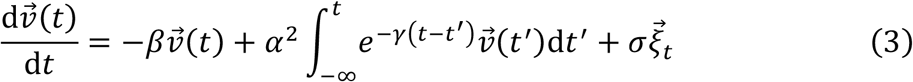

Here, 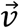 is the velocity, and 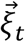 is a normalized Gaussian white noise that satisfies 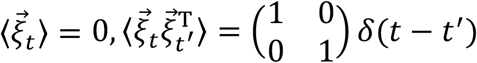, ⟨ ⟩ is an ensemble average and δ(*t*) is the delta function, *σ* is the noise strength (Selmeczi et al., 2005). By introducing

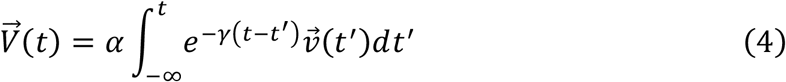

the equation of motion becomes

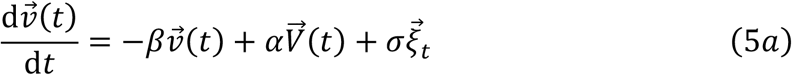

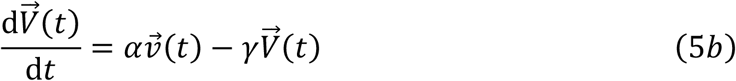

Based on the values of *T*_1_, *T*_2_, Φ_1_, Φ_2_ obtained above, we calculated the parameter values of the generalized Langevin equation (**Eqs. 5a,b**) from the analytically obtained VAC at the steady state (see **Eq. 29**).

Trajectories, the MSD and the velocity autocorrelation were obtained by numerically calculating **Eqs. 5a-b** with the parameters obtained above (**Table 1**). The individual trajectories consist of combination of persistent movement and turns (**Figure 4A**). The MSD had a slope of 1.8, which matched well with the experimental data (**Figure 4B**). The distribution of |*v*| showed a single peak that was slightly smaller compared to the experimental data (**Figure 4C**). The median was 56 μm/min in the simulation, which matched well with 60 μm/min in the experiment. The velocity autocorrelation consists of two slopes that crossed over at around 10 sec (**Figure 4D red**), which was similar to the crossover in the experimental data (**Figure 4D black**). Velocity dependence of acceleration also matched well with the experimental data (**Figure 4E**). On the other hand, the range of cell speed at which turning occurred in the simulations was somewhat broader (0 to 1.2 μm/sec) compared to the real cell (0 to 0.8 μm/sec) (**Figure 4F**). While the MSD and the VAC characteristics were well captured by the memory effect described in Eq. (2), deviation from the model became evident when comparing autocorrelation separately for the cell movement (absolute velocity 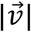) and the orientation 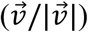 (**Supplementary Figure S3**). In the experimental data, it is only the autocorrelation of the orientation 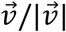 not 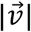 that followed double-exponential decay (**Supplementary Figure S3A, B**). In the generalized Langevin-equation, the velocity and the orientation share the same time scales, and thus the autocorrelation of both the orientation 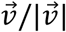 and 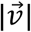 decayed with the two exponents (**Supplementary Figure S3C**,**D**).

**TABLE 1.** Parameters for the generalized Langevin equation. The experimental data were fitted with the analytical VAC (**Eq. 29**).

| | $\alpha$ [$s^{-1}$] | $\beta$ [$s^{-1}$] | $\gamma$ [$s^{-1}$] | $\sigma$ [$\mu m \cdot s^{-3/2}$] | $\sigma_x$ [$\mu m$] |
| --- | --- | --- | --- | --- | --- |
| GLE | 0.0741 | 0.116 | 0.0641 | 0.266 | 0 |
| GLE w/<br>positional<br>uncertainty | 0.0741 | 0.116 | 0.0641 | 0.266 | 0.155 |

**FIGURE 4.**
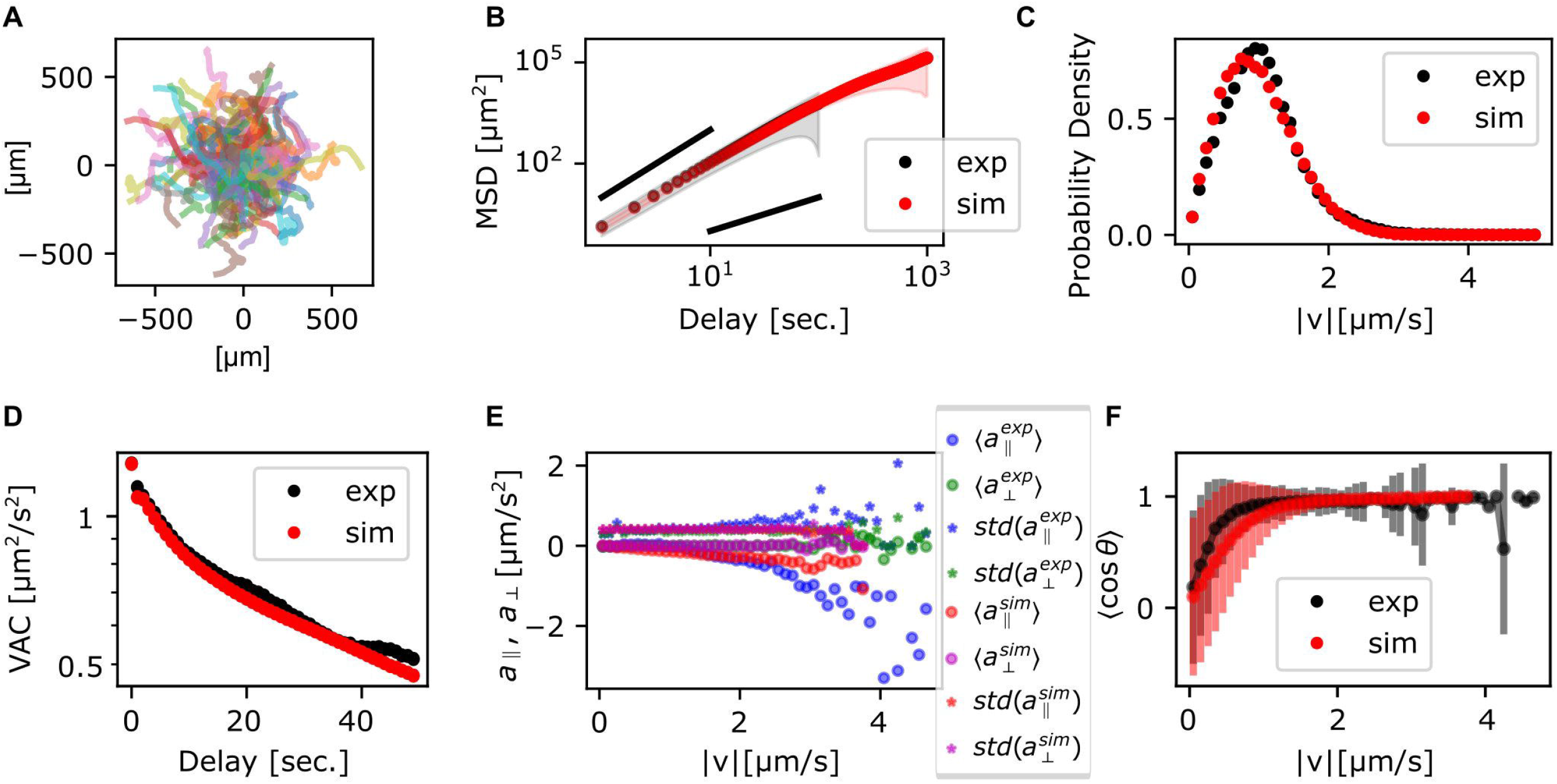
Statistics of the persistent random walker trajectories. **(A)** Simulated trajectories. **(B)** MSD; the ensemble average (circle) and the standard deviation (shade) (red). Experimental data (black) are duplicated from **Figure 3B** for comparison. **(C, D)** Probability distribution of 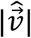 (C) and VAC (D) (red). Experimental data from **Figures 4C, D** (black) are duplicated for comparison. **(E)** Acceleration, parallel (red) and orthogonal (purple) to the velocity. The average (circle) and the standard deviation (star) for the experimental data in **Figure 3E. (F)** The average of cos *θ* (red circle) and the standard deviation (red error bar). Experimental data from **Figure 3F** (black circle and error bar) are duplicated for comparison.

Rather than pursuing extension of the particle-based formalism such as to treat the two timescales separately (Li et al., 2008; Takagi et al., 2008), we sought to more directly characterize cell reorientation by analyzing the cell shape dynamics. Based on binarized cell mask images and a boundary tracking algorithm (Nakajima et al., 2016), 500 points along the edge of cell masks were tracked in the laboratory frame for the local curvature and the normal velocity (**Figures 5A, B**; see also **Supplementary Figures S4A, C, E** for another sample). A protruding edge can be seen as a positive local-maximum in the curvature (**Figure 5A** yellow regions). The advancing front of a cell can be discerned by its positive velocity (**Figure 5B**, yellow regions), and the trailing uroid by the negative velocity (**Figure 5B**, blue regions). At the cell front, a new protrusion frequently appeared to split off from a pre-existing protrusion (**Figures 5A, B** white arrows). These appeared in the kymograph as branching positive curvature regions that propagated rearward until they were annihilated at or near the uroid (**Figure 5A** black arrows). The sequence of curvature wave dynamics represents well the shape dynamics as seen from the snapshots (**Figure 2B** white arrows). They are surprisingly similar to those obtained for *Dictyostelium* and neutrophil-like HL60 cells (Driscoll et al., 2012; Imoto et al., 2021) with a noticeable difference that splitting was more frequent and thus numerous. Occasionally, there were cases where cells transiently took dumbbell-shape (**Supplementary Figure S4B, D, E**; **Supplementary Movie S5**). In such cases, the centroid velocity orientation showed discontinuous change (**Supplementary Figure S4B**, black arrow). In the kymograph representation, a dumbbell-like cell shape appears as two or three stable curvature waves (**Supplementary Figure S4D**, black arrow). Most positions had zero velocity (**Supplementary Figure S4F**, black arrow), indicating stalling of cell shape change. These observations indicate that as the dumbbell shape appeared, the cell stopped and randomized its orientation. There were also rare cases where the cell maintained mono-polarity for an extended period of time (**Figures 5C-E, Supplementary Movie S6; see Supplementary Figure S5** for additional samples). There, new curvature waves emerged frequently and traveled fast before disappearing at the tail (**Figure 5C**). The position where curvature waves appeared always showed positive velocity, while the positions where curvature waves disappeared showed negative velocity (**Figures 5C, D**). These patterns in the kymograph correspond well with the observation that the fast curvature waves start propagating from the advancing cell front and disappear at the uroid (**Figure 5E**).

**FIGURE 5.**
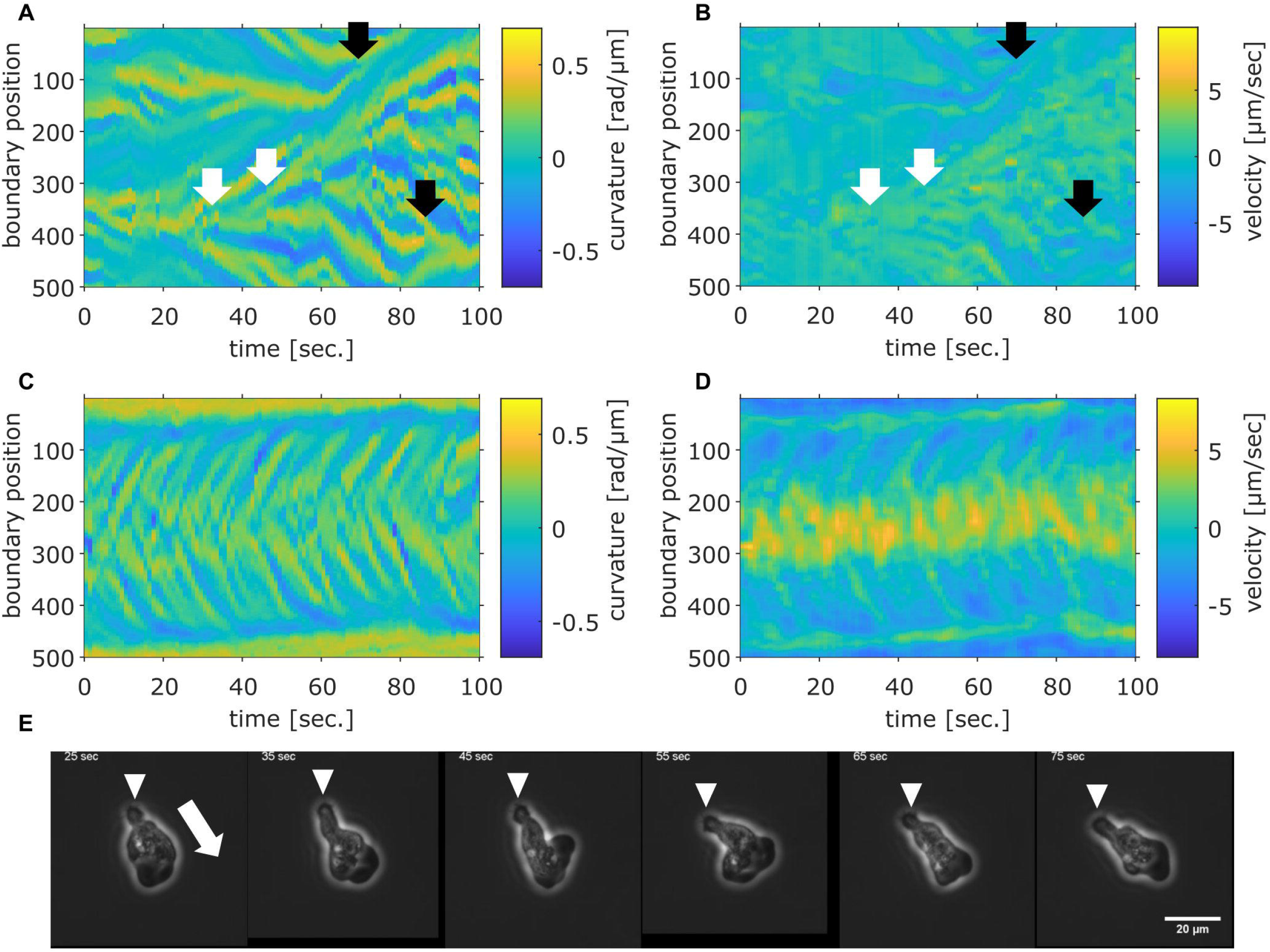
Cell boundary analysis. **(A, B)** The curvature (A) and the normal velocity (B) of the cell boundary taken from a representative cell exhibiting random walk. White arrows: splitting. Black arrows: pair annihilation. White arrows: splitting. Black arrows: pair annihilation. **(C, D)** The curvature (C) and the normal velocity (D) of the boundary taken from a cell with high persistence. **(E)** Snapshots of the cell analyzed in (C, D). The white arrow: the direction of the centroid movement. The inverted triangles mark the uroid.

A further analysis showed a close relationship between the curvature wave and the centroid movements. The protruding and the retracting membrane regions can be identified as positive curvature regions with positive (**Figure 6A**, white dots) or negative (**Figure 6A** black dots) velocity respectively. The orientation of the normal vector at the protruding region showed high correlation with the direction of centroid velocity (**Figure 6B** blue). The retracting regions oriented in the opposite directions which appeared somewhat broader in distribution (**Figure 6B** orange). To further analyze the dynamics of the curvature wave, high curvature regions (**Figure 6C** white) at each time frame were assigned as individual protrusions (**Figure 6C** green). While there were multiple protrusions in the protruding region, a dominant leading edge can be detected from identifying a single protrusion whose normal vector angle was the closest to that of the centroid velocity (**Figure 6C** magenta). Once a curvature wave became the leading edge, it remained so for about 2.8 sec as measured from its average lifetime (**Figure 6D**). Another interesting feature of the membrane extensions is that they gave birth to secondary pseudopods or were steered to other directions. The typical angular velocity associated with this dynamic was 0.1 rad/sec (**Figure 6E**). Together with the double exponential decay (**Supplementary Figure S3B**), these behaviors indicate that the centroid velocity angle by itself follows 1D persistent random walk. From experimentally obtained parameters of the leading edge lifetime (2.8 sec) and the angular velocity 0.1 rad/sec, the 1D model (see Materials and Methods, Cell Boundary Analysis section) yield decay time of 142 sec on average which matched well with the experimental data (**Figure 6F**).

**FIGURE 6.**
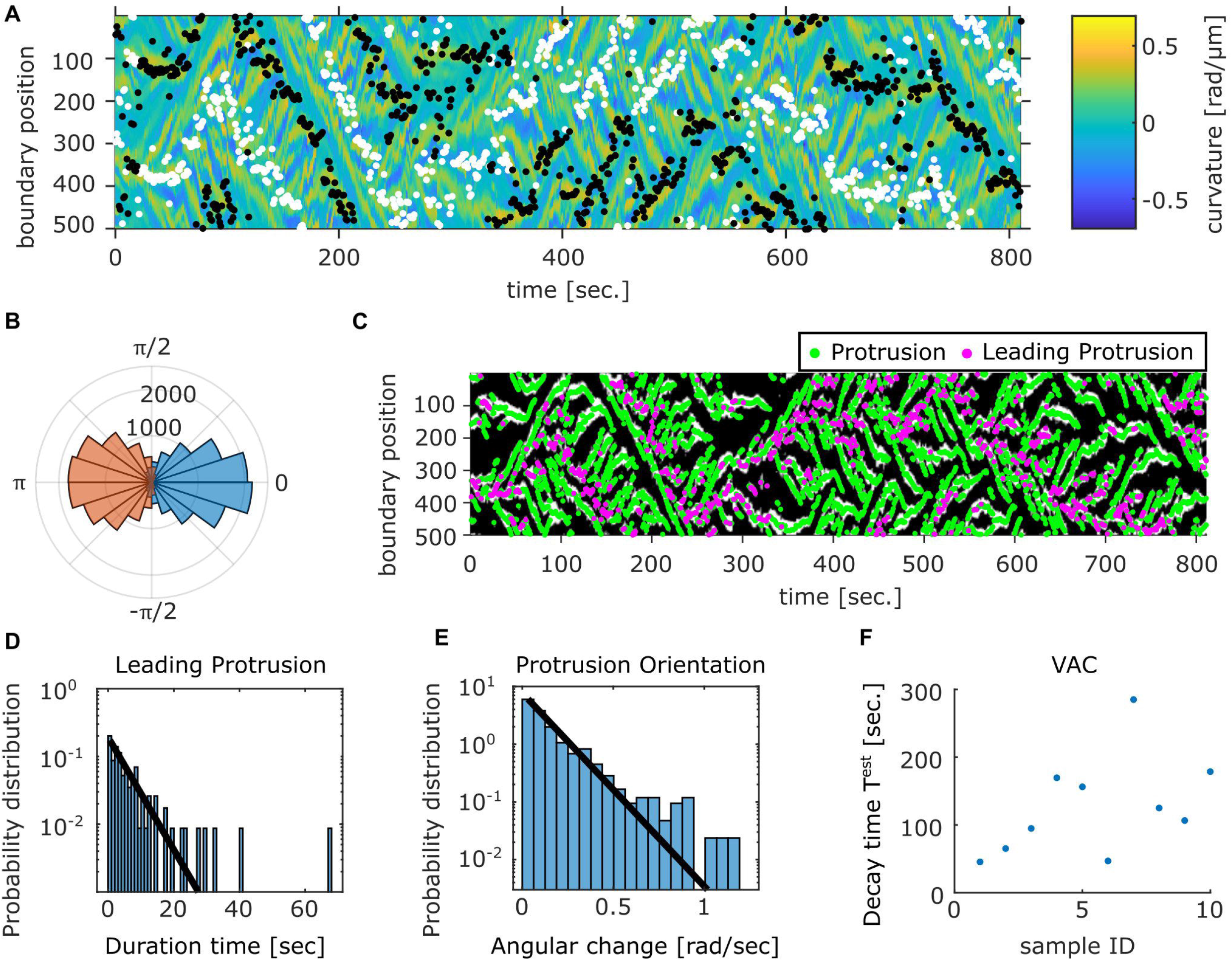
Relation between the membrane protrusions and the centroid velocity angle. (**A**) The protruding front (white) and the retracting rear (black) detected from the velocity kymograph are overlaid on top of the curvature kymograph (see Methods). (**B**) The angular histogram of the protruding front (blue) and the retracting rear (orange) relative to the cell orientation as determined by the centroid velocity. (**C**) The position of protrusive regions (‘curvature waves’; green). The region that co-extended most closely in the direction of the cell centroid motion (‘leading protrusion’; magenta). The binarized mask of the protrusion region (white) obtained from the curvature kymograph is shown in the background. (**D**) Duration time histogram of the leading protrusion (magenta in (C)). (**E**) Histogram of the angular change per unit time in the protrusion orientation (the vector normal to the cell contour at positions indicated in green in (C)). Solid lines are exponential fit to the data (D, E). (**F**) Estimated VAC decay time *T*^*est*^ for the representative data.

To obtain a quantitative morphometry, we chose by eye 21 representative mask images each for the 3 shapes; fan-shape, split and dumbbell (**Supplementary Figure S6A**) and calculated the Fourier power spectrum of the cell edge coordinates and their principal components were calculated (see Materials and Methods). We found that the first two principal components were sufficient to obtain well separated clusters that represented the shape class (**Figure 7A**). All cell masks analyzed were distributed within a confined domain in the PC1_fourier_ -PC2_fourier_ space (**Supplementary Figure S6B**). The fan-shaped data were located at a low PC1_fourier_ and high PC2_fourier_ region (**Figure 7A** circles). The split-shape were found in the low PC1_fourier_ - low PC2_fourier_ region (**Figure 7A** asterisks). The dumbbell-shape was located at high PC1_fourier_ and high PC2_fourier_ (**Figure 7A** triangles). To see what shape features the principal components represented, we reverse calculated an artificial form by obtaining Fourier spectrum from the principal component vector and the eigen vector matrix (see Methods). In brief, PC1_fourier_ indicated the aspect ratio i.e. elongation, PC2_fourier_ the head width, PC3_fourier_ the rear steepness (**Supplementary Figure S6C**). Here, the main contribution to PC1 were from the wave number 1 and -1 with coefficients of 0.68 and 0.73. For PC2, the contribution from wave number 1, -1, 2 and 3 was 0.62, -0.59, -0.49, -0.12, respectively. Contribution from other modes was small with coefficients less than 0.03.

**FIGURE 7.**
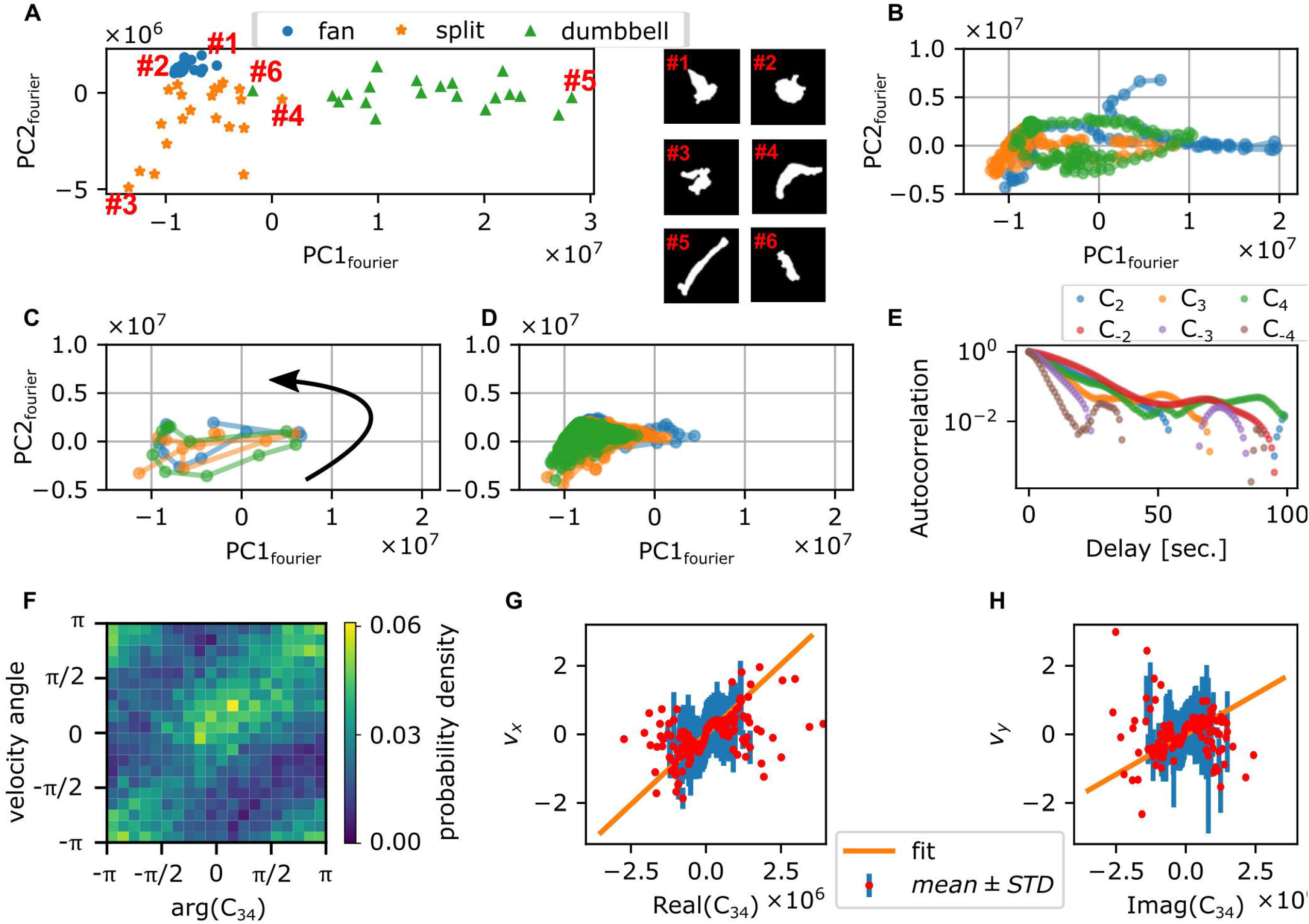
Fourier analysis of the cell contour. (**A**) Principal component space (PC1_fourier_, PC2_fourier_) obtained from 63 manually selected binarized snapshots (left panel). Representative cell masks (right panels). (**B**) Time series in PC1_fourier_-PC2_fourier_ space from 3 representative timelapse sequences (colors). (**C**) Time evolution of PC1_fourier_ and PC2_fourier_ of 10 sec around a large turn that involves transition to the dumbbell shape (3 representative events; colors). Black arrow indicates the direction of time evolution. (**D**) Time evolution of PC1_fourier_ PC2_fourier_ during persistent migration. Colors indicate different time series (duration: 269 sec (blue), 1039 sec (orange), or 3600 sec (green)). (**E**) Autocorrelation of *C*_n_ (n = ±2, ±3, ±4). The decay rate: 12.2 (*C*_2_), 18.0 (*C*_-2_), 8.9 (*C*_3_), 7.6 (*C*_-3_), 10.6 (*C*_4_), and 4.4 (*C*_-4_) seconds. (**F**) Distribution of angles of centroid velocity and *C*_34_. (**G, H**) The x- (**G**) and y-components (**H**) of centroid velocity plotted against real (**G**) and imaginary (**H**) parts of *C*_34_. Red circles and blue bars indicate average and standard deviation of centroid velocity binned with the value of *C*_34_. Orange lines indicate the result of fitting with linear proportionality.

How the cell shape changed during turning can be analyzed by tracking the time sequence in the PC1_fourier_ - PC2_fourier_ space. **Figure 7B** shows three independent samples of re-orienting cells. Here, cells were mainly located in the negative PC1_fourier_ region with occasional visits to the positive PC1_fourier_ region. This is consistent with the above observation that cells took fan-or branched-shape (negative PC1_fourier_) in addition to rare occurrence of dumbbell-shape (positive PC1_fourier_). **Figure 7C** shows three independent samples of the dumbbell-shape forming cells. The counter-clockwise circular trajectories in the PC1_fourier_ - PC2_fourier_ space signify a transition from the fan-shape to splitting then to the dumbbell-shape. On the other hand, **Figure 7D** shows three independent trajectories that remained in the negative PC1 region for extended period of time. These cells at least during the time window of observation fluctuated between the fan-shape and the bifurcating fingers. There was no clear relationship between the morphometry state (PC1_fourier_, PC2_fourier_) and the cell speed **(Supplementary Figures S7A, B**). There was, however, negative correlation between the centroid speed and the rate of state transition *d*{PC1_fourier_}/*dt* but not with *d*{PC2_fourier_}/*dt* (**Supplementary Figures S7C, D**). As the former relation was seen in the negative direction *d*{PC1_fourier_}/*dt* <0, it signifies that cells accelerate when recovering from dumbbell-shape.

Besides the rate of state transition in the principal components, there should be a direct relationship between the Fourier components *C*_n_ themselves and the centroid movement. Autocorrelation analysis showed that the decay rates for *C*_-3_, *C*_3_ and *C*_4_ were 7.6, 8.9, and 10.6 second, respectively (**Figure 7E**) and thus matched most closely to the short decay time of VAC. As for the centroid velocity itself, according to the deformation tensor-based theory of cell movement (Ohta et al., 2016), it should be proportional to *C*_*nm*_ ≡ *Ċ*_− *n*_*C*_m_− *C*_− *n*_*Ċ*_*m*_ − *C*_− *n*_*Ċ*_*m*_ where *n* + *m* = 1. More specifically, *C*_*nm*_ is a complex number whose absolute value |*C*_*nm*_| and the angle arg(*C*_*nm*_) are expected to be proportional to the speed and the velocity angle of the centroid respectively. In NIH3T3 cells, it has been shown that velocity is proportional to the elongation *C*_-2_ and triangular *C*_3_ modes of deformation multiplied by their time derivatives; i.e. *C*_23_ = *Ċ*_− 2_*C*_3_− *C*_− 2_*Ċ*_3_ (Ebata et al., 2018). However, in *Naegleria gruberi*, we found little correlation between *C*_23_ and the centroid velocity (**Supplementary Figures S7E-G**). Instead, we found that it was *C*_34_ = *Ċ*_− 3_*C*_4_ − *C*_− 3_ *Ċ*_4_ that correlated highly with the centroid velocity angle (**Figure 7F**) and x- and the y- component of the centroid velocity (**Figures 7G, H**). The difference between *Naegleria* and NIH3T3 may be attributed to the fact that *Naegleria* has many pseudopods that are complex in shape as analyzed below.

To further investigate the cell shape characteristics, we employed a convolutional neural network that was previously trained to classify cell shapes based on similarity to *Dictyostelium*-like, HL60-like, or fish keratocyte-like shapes (Imoto et al. 2021). While the method is not suited to track shape change over time due to discrete change in the morphometry space that is sometimes introduced by uncertainty in the cell orientation during mask alignment, it has an advantage of providing an objective morphometry that is independent of known feature basis. On average, Naegleria was classified as *Dictyostelium*-like (high PC1_cnn_, low PC2_cnn_) (**Figure 8A**). This was natural as it has been shown to pick up branching shapes that are elongated overall in the migrating direction (Imoto et al., 2021). We noticed substantial variability, however, in the individual cell shapes (**Figure 8B; black)** that exceeded those normally observed in *Dictysotelium* (Figure 8B; green). Shapes that deviated in the PC1_cnn_ direction were mapped to dumbbell-like domain in the Fourier descriptor-based morphometry (**Figure 8C; orange**). Those that deviated towards low PC1_cnn_ were mapped to the domain that showed numerous pseudopods (**Figure 8D; orange**). Datapoints that fell at or near the HL60-like domain (low PC1_cnn_ low PC2_cnn_) were mostly fan-like (**Figure 8C**; blue, **Figure 8D**; blue) and their occurrence per timeseries showed positive correlation with the MSD exponent (**Figure 8E**). This is consistent with the notion that more mono-polarized the cells are, the more ballistic the cell trajectories become.

**FIGURE 8.**
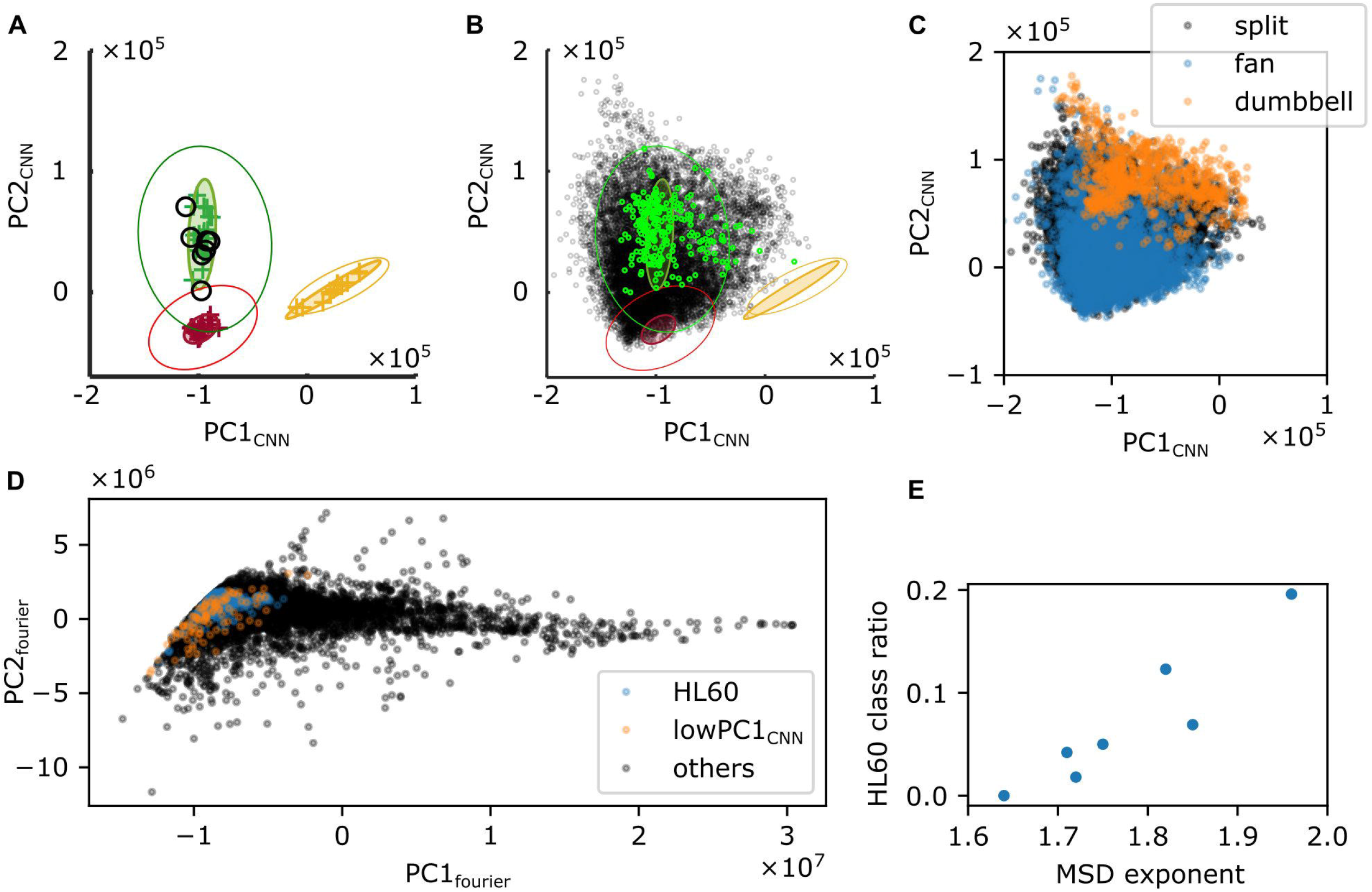
Shape analysis by a CNN-based classifier. (**A, B**) Time-average (A) or snapshot (B) of PC1_CNN_ and PC2_CNN_ from N. gruberi images (black) were superimposed on the PC1_CNN_ -PC2_CNN_ phase space of *D. discoideum* (green), HL60 (red), and fish keratocyte (yellow). (**C**) Snapshot data in (B; black) was classified into split (black), fan (blue) or dumbbell (orange) based on PC1_fourier_ and PC2_fourier_. (**D**) PC1_fourier_ and PC2_fourier_ of HL60-like (blue), cells with PC1_CNN_ lower tha n -1.6×10^6^ (orange), or the other cells (black) classified with CNN. (**E**) Ratio of frames whose shape was classified as HL60 in deep leaning-based classification.

## DISCUSSION

In this report, we analyzed movements of *N. gruberi* cells by quantifying their speed, directionality, and shape change. The locomotive speed of *N. gruberi* cells was around 60 μm/min, which is similar to that reported in early literatures (King et al., 1981; Thong and Ferrante, 1986). It is substantially larger in magnitude compared to that of fibroblast ∼ 0.4 -1.0 um/min (Welf et al., 2012; Passucci et al., 2019), and even larger compared to fast migrating cells such as vegetative *Dictyostelium* 5 μm/min (Li et al., 2008), and neutrophils 17 μm/min (Hartman et al., 1994). Despite the large speed difference, we found common features between *N. gruberi* and other cell types whose random motility have previously been characterized. The exponent of MSD was approximately 1.8 meaning that the random walk is non-Fickian at least at surface. Similar exponent is known in MDCK cells (Dieterich et al., 2008), A549 cancer cells (Kwon et al., 2019) hematopoietic progenitor cells (Partridge et al., 2022), and T cells (Jerison and Quake, 2020). Of particular note is that the time-scale where such exponent was observed for *Naegleria* was about 10 to 100 sec which is within the order of magnitude required for a cell to move one cell-body length. This seems also to be the case for MDCK cells where the exponent of 1.8 was observed at much longer time-scale of 4 to 20 min with corresponding length scale 4 μm to 20 μm. All in all, our data combined with the observations above from earlier literatures suggest that the time scale at which cells move in a straight line is the major determinant of cells’ displacement.

The other common feature found in this study was the presence of two characteristic decay time in the velocity autocorrelation (Selmeczi et al., 2008). For *Naegleria*, these were *T*_1_= 6 and *T*_2_ = 90 sec, which are in the same order of magnitude as that of *Dictyostelium* in the vegetative (*T*_1_ = 5.2 and *T*_2_ = 228 sec) and the starved (*T*_1_= 11 and *T*_2_ = 108 sec) states (Li et al., 2008). Although equivalent measurements have not been documented for neutrophils, their cell shape changes had typical time scale of 8 sec (Hartman et al., 1994) and the persistence time during chemotaxis was 103 sec (Itakura et al., 2013; Haastert, 2021). From the MSD measurement, the persistent time of *Dictyostelium* and neutrophil-like HL60 were 151 sec and 278 sec, respectively (Imoto et al., 2021). Interestingly, VAC of Human keratinocyte-like cells (HeCaT) whose speed was much slower (0.18 μm/min) could also be fit with double exponential (*T*_1_= 76 sec and *T*_2_ = 860 sec; (Selmeczi et al., 2005)). The characteristic time scale of around 10 sec was attributed to the time scale of actin polymerization in the protruding pseudopodia (Haastert, 2021). However, the pseudopod lifetime in *N. gruberi* was rather long; about 15 to 50 sec. The discrepancy may be attributed to the sister pseudopods that formed from the main pseudopods which were not analyzed in our manual tracking. In some cases, the pseudopod itself also appeared to bend in one direction. In support of this notion, the autocorrelation of *C*_-3_ and *C*_4_ had decay time of 7∼10 sec which matched well with the first decay time of VAC.

On the other hand, the second decay time of VAC (90 sec; **Figure 3D**) was close to the timescale of directional persistence i.e. ‘run’ phase estimated from the curvature wave dynamics (142 sec; **Figure 6F**). As for the average cell speed, we found strong correlation between the centroid velocity with the coupling of deformation modes *C*_-3_ and *C*_4_, instead of *C*_-2_ and *C*_3_. This suggests that the orientation *of Naegleria gruberi* cells depends not on the main membrane protrusion but on their sister sub-structures. A further pseudopod-level analysis at finer time-scale is required to clarify the relation between the deformation modes and the branching pseudopods. The rare cells with high persistency did not take high PC1_fourier_ value (**Figure 7D**) which was opposite of *Dictyostelium* (Tweedy et al., 2013). This likely stems from the fact that the elongated form in *Naegleria* was usually dumbbell-shaped which occurs when cells stall and reorient.

The splitting pseudopod dynamics may entail a mechanism similar to those found in amoebozoa and metazoan that involves excitable and oscillatory generation of dendritic actin meshworks (Huang et al., 2013). The presence of local inhibitor of pseudopod formation in neutrophils and *Dictyostelium* (Xu et al., 2017) and potential lack of such in *Naegleria* may underlie the difference in the number of pseudopods. Alternatively, there may be local reduction in the actin cortex that are stochastic in nature. Although protrusions observed under our culture conditions did not appear as blebs, flow of cytosol towards the membrane during extension of a protrusion suggests local pressure release. The extension of protrusion triggered by concentration of pressure on a certain point of the surface is known as viscous fingering. The movement speed of *N. gruberi* was 5 times as large as that of neutrophils and *Dictysotelium*, but closer to that known for fragments of *Physarum* which was also known for marked cytoplasmic streaming (Rieu et al., 2015) and persistent random walk (Rodiek and Hauser, 2015). Such high velocity and potential interface instability may also underlie the observed branching of pseudopods. Another unique shape feature was the dumbbell-like cell shape. According to a recent theoretical model of a lamellipodia-based dynamics, a similar “two-arc shape” appear when the protrusive force is high (Sadhu et al., 2023). The dumbbell-shape may thus be a prevalent shape feature that were heretofore overlooked due to peculiarity of the model cells. Indeed, a similar dumbbell-shape has been reported in fragmented *Physarum polycephalum* (Rieu et al., 2015).

In the *E. coli* run-and-tumble, the underlying biochemical network has been proposed to be optimally designed to extract binary information in a noisy environment (Nakamura and Kobayashi, 2021). Some bacterial species make use of multiple run modes that differ in how they are modulated in the presence of chemoattractants (Alirezaeizanjani et al., 2020) suggesting diversity and depth at which random walk strategies are likely employed in prokaryotes. In *Dictyostelium* amoebae, the run length increases under starvation (Haastert and Bosgraaf, 2009) which may be related to their foraging strategy. In immune cells, high correlation between cell speed and persistence is thought to underlie their search efficiency in vivo (Shaebani et al., 2020, 2022). Cancer cells shows persistent random walk in the metastatic state while weakly persistent in non-metastatic state (Huda et al., 2018). Although chemoattractant of *Naegleria gruberi* is so far unknown, in *Naegleria fowleri*, formylated peptides are known to act as chemokine (Marciano-Cabral and Cline, 1987) meaning that it enhances cell polarity and movement in the absence of gradient. Cell-cell variability in such response may explain how a minority of *Naegleria gruberi* cells under our experimental condition showed persistent monopolarity. In future works, it should be informative to study how the behaviors quantified in this work are modulated by chemotactic and chemokinetic factors and how they are related to exploratory and invasive strategies.

## MATERIALS AND METHODS

### Cell culture

*Naegleria gruberi* strain NEG-M was obtained from American Type Culture Collection (ATCC 30224). For routine cell propagation, small bits of frozen stock were scraped off using a sharp needle onto a fresh lawn of *Klebsilella aerogenes* on a NM agar plate (Peptone, Dextrose, K_2_HPO_4_, KH_2_PO4, 2% bactoagar)(Fulton, 1970). The two-member culture plate was incubated at 30 °C for a few days until cleared plaques appeared. To start axenic culture, growing cells were picked from the edge of a plaque and suspended in milliQ water. 10 μL of the cell suspension was added to 25 mL modified HL5 media (Fulton, 1970) supplemented with 40 ng/mL vitamin B12 and 80 ng/mL folic acid, 10% fetal bovine serum (FBS, Sigma 172012) and 1% Penicillin-Streptomycin (Gibco) in a 75 cm^2^ canted-neck plastic flask (Corning 431464U). Cells were allowed to attach to the bottom of the flask and incubated at 30 °C for 3 days before harvesting for imaging.

### Time-lapse imaging

Axenic growing cells were dislodged from the flask bottom by gentle agitation. Cells were pelleted by centrifugation at 7 × 10^2^ G for 3 min and resuspended in fresh HL5 media. The medium contains 5 mM KH_2_PO_4_ /Na_2_HPO buffer and thus provides required electrolyte (King et al., 1979) for optimal migration. The cell density was adjusted to 3.3 × 10^2^ cells/mL for the observations. 3mL of the cell suspension was plated on a 35 mm glass bottom dish (No.0 20 mm hole diameter, MatTek). The plate was set to the stage of an inverted microscope (IX81, Olympus) equipped with either a thermal plate or a closed stage-top incubator set to 30 °C and kept still for 30 min before starting time-lapse image acquisition. All image acquisition was performed at 30 ° C.

Phase contrast images were obtained by 40x (LUCPLFLN) objective lens and a sCMOS camera (Prime 95B, Photometrics). To track a target cell at multiple non-overlapping fields of view (FOV), Micromanager software with a custom written plugin was employed. Timelapse images were obtained from 2 or 3 positions at an interval of 1 second for up to 1 hour. Each position was chosen so that initially only a single cell at the center existed in the entire field of view. In between each image acquisition, the cell centroid was calculated from a mask obtained by applying the ‘Make Binary’ function in ImageJ to the most recent image. The automated stage was then recentered to cancel out the centroid displacement.

### Analysis

#### Characteristics of cellular trajectories

Binary masks from timelapse images were prepared using LABKIT (Arzt et al., 2022). Trajectories of cell centroid were extracted from the mask images using the ImageJ plugin TrackMate (Ershov et al., 2022). Trajectories from simulations of the generalized Langevin equation (**Eqs. 4-6**) were sampled at every 1 second using the Euler-Maruyama method with 2-milisecond interval with open library TorchSDE (Li et al., 2020; Kidger et al., 2021). Velocity 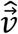 and acceleration 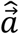 were calculated from the difference in the sampled positions 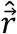 at time *t*_n_ = *nδt* with an interval *δt*:

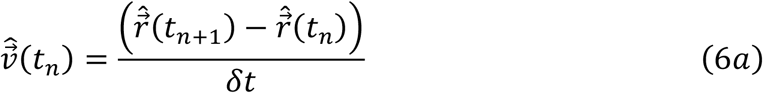

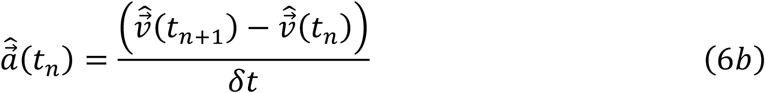

MSD *msd*(mδt), probability distribution of speed *p*(*v*), velocity autocorrelation *vac*(mδ*t*), mean and standard deviation of acceleration conditional on speed 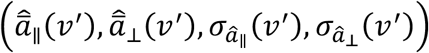, and conditional-averaged strength of turning ⟨cos *θ*⟩(*v*′) were calculated from the trajectories for both the experiment and simulation data according to

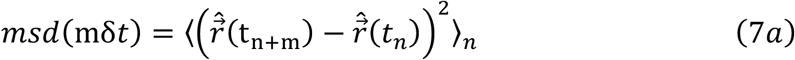

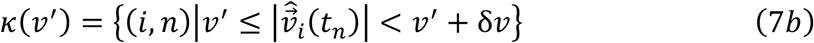

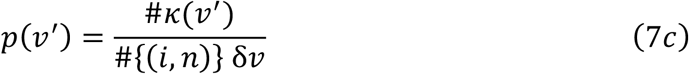

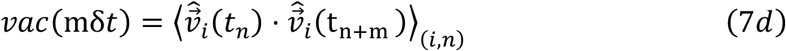

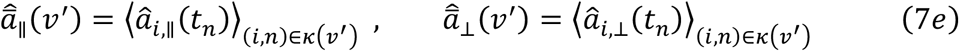

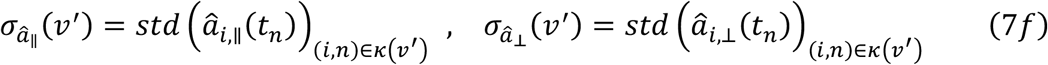

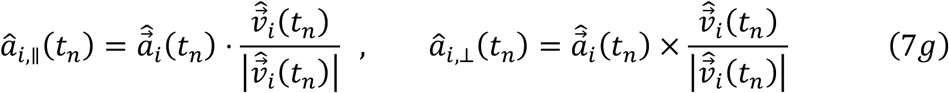

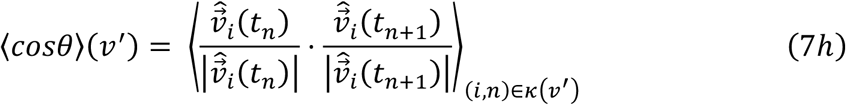

where ⟨ ⟩_*X*_ is the average over *X*, subscript *i* indicates the *i*-th trajectory, # is the number of items in the following set {}, *std* denotes the unbiased standard deviation, and δ*v* = 0.1 μm/sec is the bin width. Additionally, we checked the detail of the time evolution of velocity by calculating the autocorrelation of the magnitude and the angle:

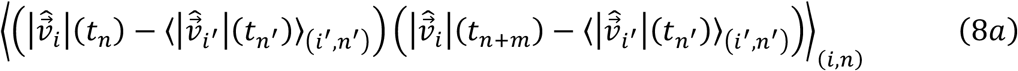

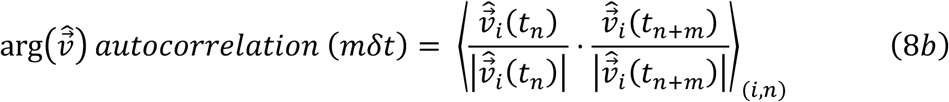

#### Velocity distribution

We fit a Gaussian distribution to both *v*_*x*_ and *v*_*y*_ to determine the standard deviation *σ*_*G*_. For 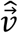 that follows 2-dimensional Gaussian distribution, the distribution of 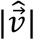 is readily derived from the chi-square distribution with 2 degrees of freedom where the square of 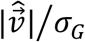 follows:

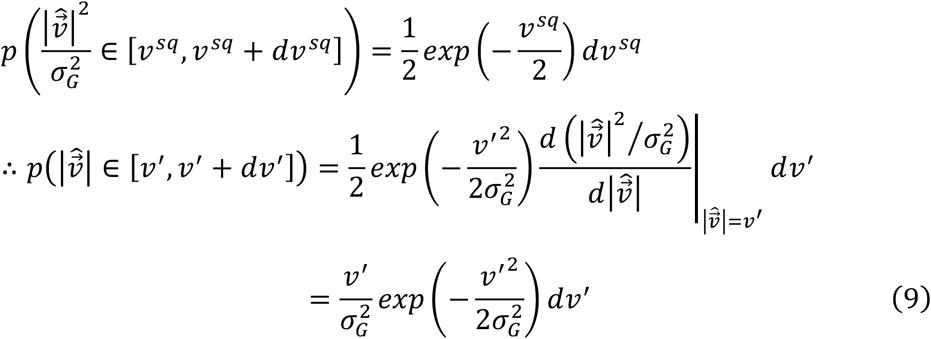

To note, the peak of the above distribution is located at 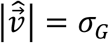.

#### Fitting VAC

To fit the experimental data with the generalized Langevin equation (**Eqs. 5a,b**), we employed the analytical solution for the velocity autocorrelation *vac*^*ss*^. For the observed velocity 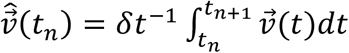

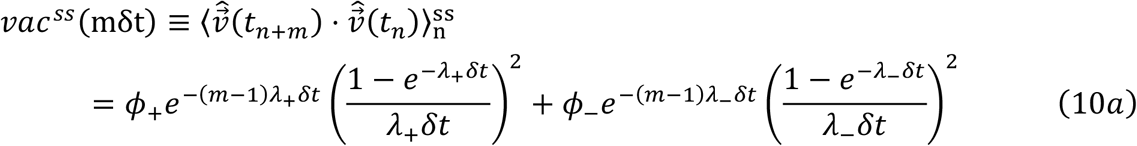

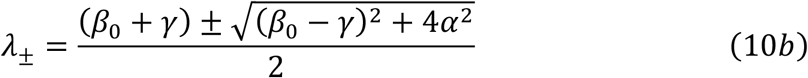

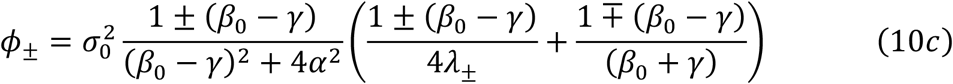

Optimal values of *α, β*_0_, *γ, σ*_0_ were obtained by minimizing the mean square error between *vac*^*ss*^(mδ*t*) and *vac*^*exp*^(mδ*t*).

#### Positional uncertainty

Parameters in **Table 1** were obtained by fitting VAC at *τ* ≥ 2 *sec*. As for the simulation only with generalized Langevin equations (**Eqs. 5a,b**), VAC matched poorly for the shortest time interval of our data (*τ* = 0 and 1 sec) due to measurement uncertainty arising from finite time step and spatial resolution of the observation. Because acceleration was also defined as the velocity difference in this time interval, the magnitude of acceleration in the simulations was off by one order of magnitude from the real cell data. We emulated these effects in the simulations by including white Gaussian noise with the observed standard deviation *σ*_*X*_ (see Methods, **Table 1**). The value of VAC changed only at the shortest time window of (*τ* = 0 and 1 sec) by this correction.

To represent positional uncertainty, we incorporated additive noise in the model so that

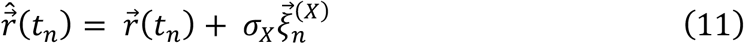

where 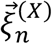is white gaussian noise which satisfies 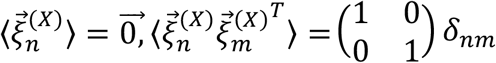 where *δ*_*nm*_ is the Kronecker delta, and thus independent of all the other variables. *σ*_*X*_ is the strength of the positional noise, 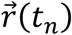 is the position sampled at time *t*_*n*_, calculated by integrating 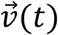 in time according to **Eqs. 5a,b**. The observed velocity 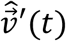 used in the analysis is defined as follows:

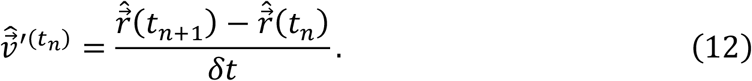

Due to the positional noise, the analytical solution of the velocity autocorrelation at steady state becomes

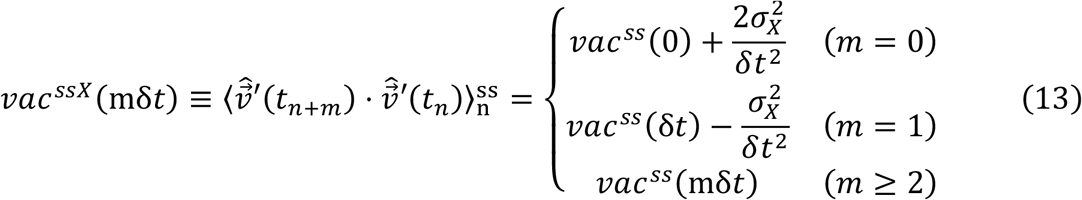

The optimal values of *α, β*_0_, γ, *σ*_0_, *σ*_*X*_ were obtained by minimizing mean square error between *vac*^*ssX*^(mδ*t*) and *vac*^*exp*^(mδ*t*).

#### Cell Boundary analysis

A MATLAB code for the active contour method (Driscoll et al., 2012) - BoundaryTrack (Nakajima et al., 2016; Fujimori et al., 2019) was used to plot kymographs of the curvature and protrusion velocity of the cell binary mask contour. In brief, the kymographs show time-evolution of curvature or normal vector-projected velocity on the contour. The angle of normal vector was also obtained using this code.

##### 1) Comparing the protrusion velocity and the cell centroid velocity

To detect the forward region of the cell, the *i* -th boundary point at time *t* in the velocity kymograph {*u*_*i*_(*t*)}_*i*=1,…,500_ were smoothed by fitting the velocity values at boundary points in each time with the following joint function:

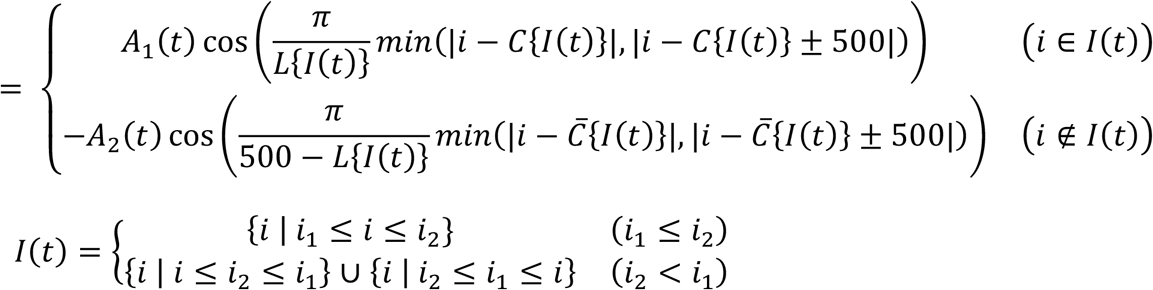

where *A*_1_, *A*_2_ ≥ 0, *I*(*t*) is a continuous front region bounded by two ends *i*_1_ (*t*) and *i*_2_ (*t*). The center *C*{*I*(*t*)}, length *L*{*I*(*t*)}, the center of rear region 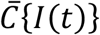 were defined in the coordinate with the periodic boundary condition, as follows:

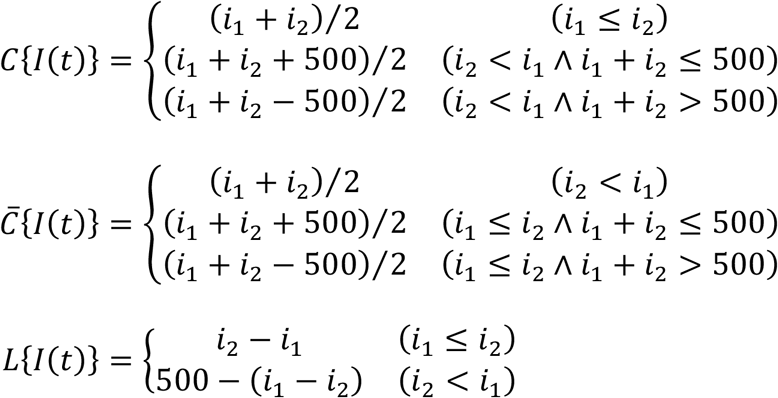

To investigate the relation of front or rear region with the direction of cell centroid velocity, we calculated the angle difference between the normal vector at the center of front or rear region and the centroid velocity.

##### 2) Curvature wave tracking and the leading edge detection

To track the curvature waves, we first detected protrusive regions as follows. Depending on the curvature {*c*_*i*_(*t*)}_*i*=1,…,500_, position #i in the curvature kymograph were classified as either ‘protrusive’ (*c*_*i*_(*t*) > *c*^(2)^), ‘flat’ (*c*^(1)^ < *c*_*i*_(*t*) ≤ *c*^(2)^) or ‘caved’ (*c*_*i*_(*t*) ≤ *c*^(1)^) where the thresholds *c*^(1)^, *c*^(2)^ were obtained by the Otsu’s method. At each time point *t*, continuous protrusive regions (j = 1, 2, 3…) were defined as set 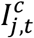 of neighboring protrusive boundary points i between two ends 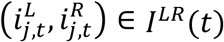:

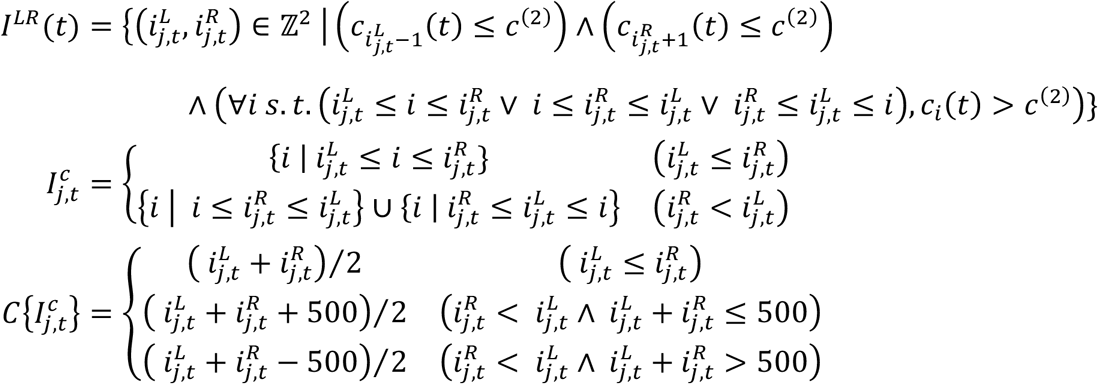

where 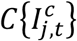 is the center of *j*-th protrusive region.

Next, we traced the curvature waves by linking the *j*-th fragment at frame *t* and the *j*’-th fragment at frame *t* + 1 if 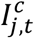 and 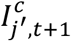 have overlapping points. Thus, the set of linked fragments *J*^*c*^ was defined as follows:

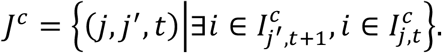

From each pair of the linked fragments (*j, j*^′^, *t*) ∈ *J*^*c*^, we obtained the angular velocity 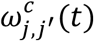 of a protruding region as follows:

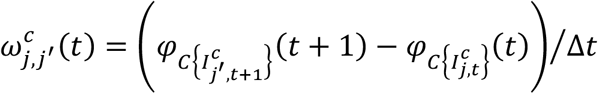

where *φ*_*i*_(*t*) is the angle of the normal vector at point *i* at time *t*. The representative angular velocity *ω*^*c*^ were obtained by fitting the histogram of 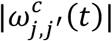 to an exponential distribution for all the linked fragments.

To investigate the relation between the curvature wave and the centroid velocity angle, we selected a single dominant wave *j*^*d*^(*t*) whose angle of normal vector 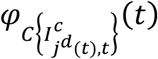 was closest to that of the centroid velocity at time *t*. The lifetime 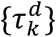 of the leading edge was measured by calculating the time window during which the leading edge was assigned to a particular curvature wave. To this end, we computed the time interval between the time points 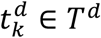 at which 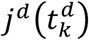 become un-linked to the dominant wave at the next time frame 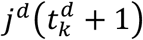:

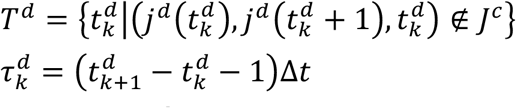

where the index *k*is given so that 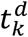 is listed in the ascending order, i.e., 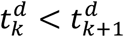 for all integer *k*. We fit a histogram of 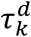 for all the linked fragments with exponential distribution to obtain the typical duration time of driving wave *τ*^*d*^.

##### 3) Estimating the time scale of centroid velocity autocorrelation

The angular velocity ω^*c*^ and the duration time *τ*^*d*^ obtained above were used to estimate the autocorrelation of the angle of cell centroid velocity *ψ*(*t*). The time evolution of *ψ*(*t*) was modeled as 1D persistent random walk with time scale *τ*^*d*^ and step size ω^*c*^*τ*^*d*^. Then the probability distribution of the angle difference Δ*ψ* ∈ (−∞, ∞) is:

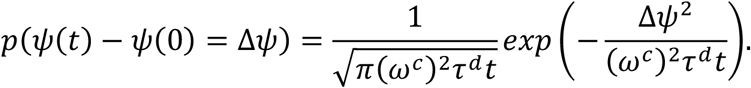

Therefore, the autocorrelation AC(t) is:

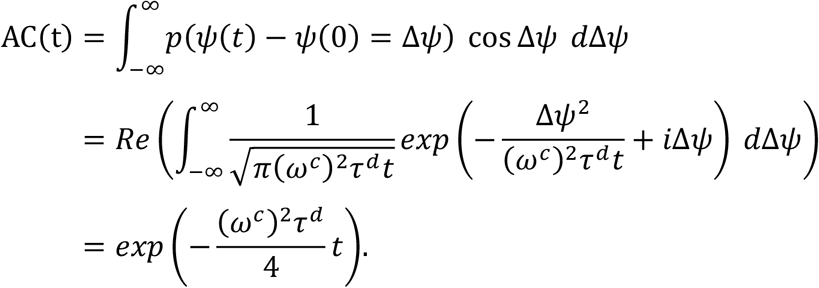

Thus, the estimated decay time of the autocorrelation is *T*^*est*^ = 4/(ω^*c*^)^2^*τ*^*d*^.

#### Cell morphology analysis

##### 1) Fourier-based shape analysis

To quantify cell shape, we calculated the elliptic Fourier descriptor (Kuhl and Giardina, 1982). First, we extracted the outline of cell binary mask with a homemade code according to (Nakajima et al., 2016; Fujimori et al., 2019). The periphery of a cell mask Γ was defined as a folded line parametrized with length 0 ≤ *ℓ* < L connecting the pixels 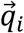 on the edge, where each pixel *i* has pixel *i* − 1 and *i* + 1 in its 4 nearest neighbor pixels:

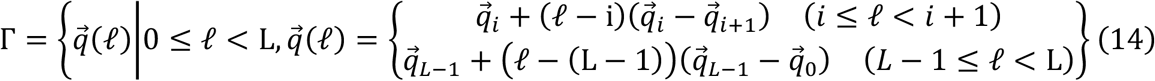

Next, the polygonal outline was converted to 160 equally spaced points 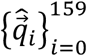 on a relative position on Γ:

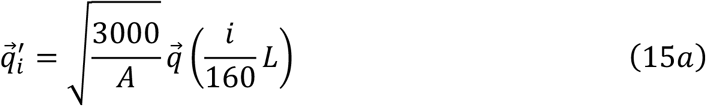

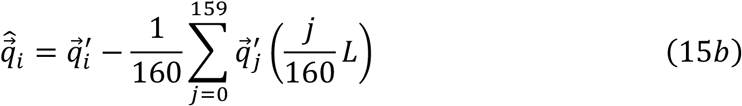

which is rescaled according to the total number of pixels *A* in the mask, and the coordinate was set so that the origin is at the cell centroid.

The elliptic Fourier descriptor was calculated by taking the discrete Fourier transformation of 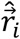 with wave number *k*:

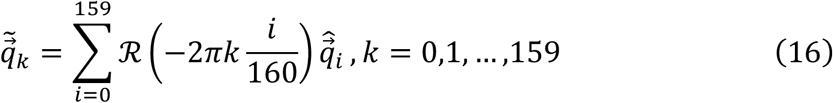

where ℛ (⋅) is a rotational matrix. Its power spectrum 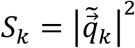 was calculated. *C*_*n*_ and *C*_−n_ are complex number equivalents of 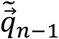 and 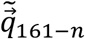.

##### 2) Fourier descriptor PCA

We calculated principal component vectors from the representative dataset containing 63 snapshots. From the power spectrum vector 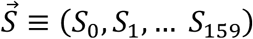 for each mask in the representative dataset, averaged power spectrum vector 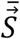 and the covariance matrix η = (η *kl*)_*k,l*=0,1,…,159_

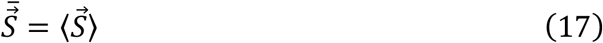

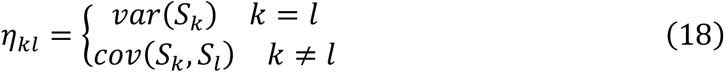

were calculated, where *var* and *cov* denotes the variance and covariance. The *m*-th eigenvalue and eigenvector 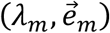 of matrix η were defined so that the conditions *λ*_1_ ≥ *λ*_2_ ≥ ⋯ ≥ *λ*_160_ and 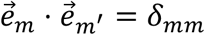_′_ are met. To note, thus obtained values of 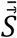,, and 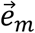 were used to analyze all the data. Using the eigenvectors, the *m*-th principal component

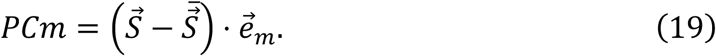

was calculated for each power spectrum vector of mask.

To characterize cell shape change dynamics, we calculated autocorrelation *AC*^*PC*^ of PC1 and PC2 values. Using the PC values of cell *i* at time *t*_*n*_, *AC*^*PC*^ is:

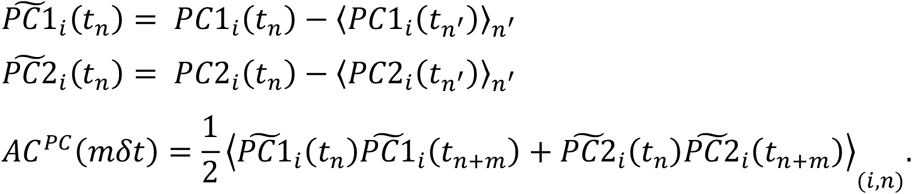

To restore the shape of cell from a set of principal components (*PC*1, *PC*2, …, *PC*160), 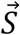 and 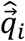 were sequentially calculated:

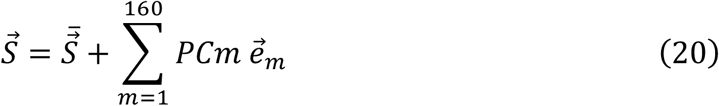

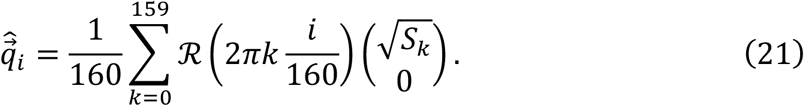

The pixels 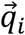 included in the edge were obtained by rounding off 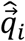. To show the recovered edge as an image, we made a binary image which has white color only on the pixels 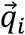.

##### 3) CNN-based shape analysis

CNN-based PCA and classification were performed based on the morphometry obtained previously (Imoto et al, 2021). In brief, each snapshot image of *N. gruberi* was input to the pre-trained CNN, and the morphology features were obtained as output. The principal components of these features were calculated using the PCA parameters obtained in (Imoto et al, 2021). The time average of the principal components was taken from all the frames in each time series. According to the morphology features, each snapshot was classified into three morphology classes: *Dictyostelium-like*, HL60-like, and fish keratocyte-like. Since only two snapshots were classified as keratocyte-like, we conduced the further analysis on *Dictyostelium-like*, HL60-like classes. The HL60 class ratio was calculated for each timeseries, as the number of snapshots classified as HL60 divided by the total number of snapshots in the timeseries.

### Analytical solution of VAC at steady state without positional noise

First, we define VAC as an ensemble-averaged inner product of true velocities at two timepoints:

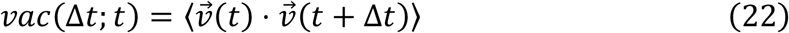

To obtain the dynamics of thus defined VAC, 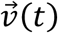 can be obtained as itô-integral of generalized Langevin equation with 2-dimensional Brownian motion 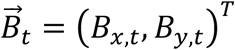 :

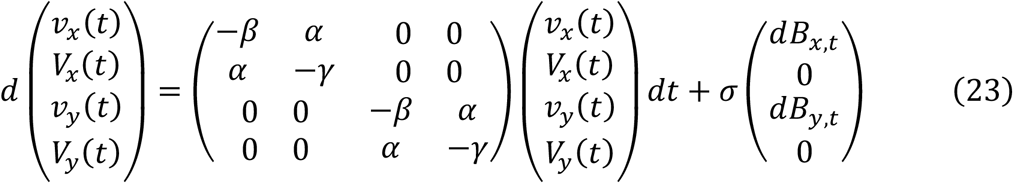

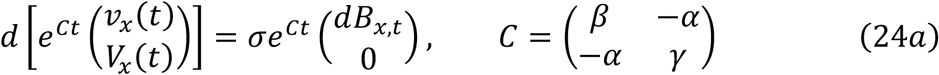

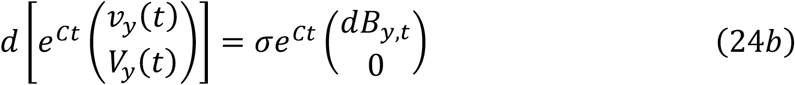

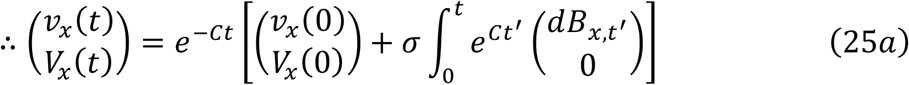

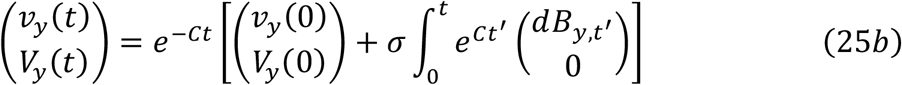

Especially, the velocity can be calculated from the eigenvalues *λ*_±_ defined above and corresponding eigenvectors 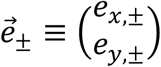 of *C* with 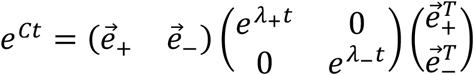:

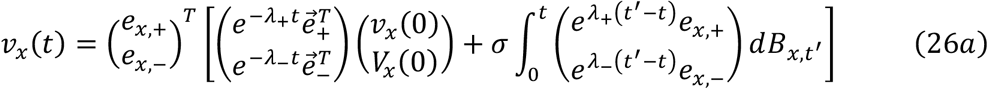

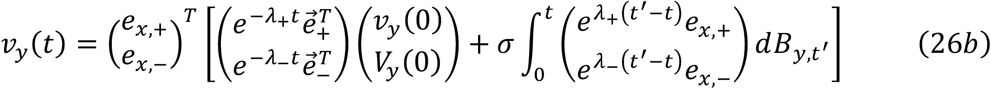

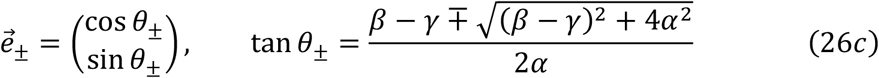

Using the representation of 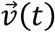 above and the with property of Brownian motion 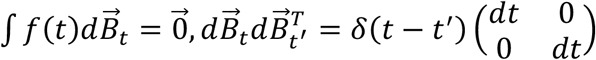,VAC is:

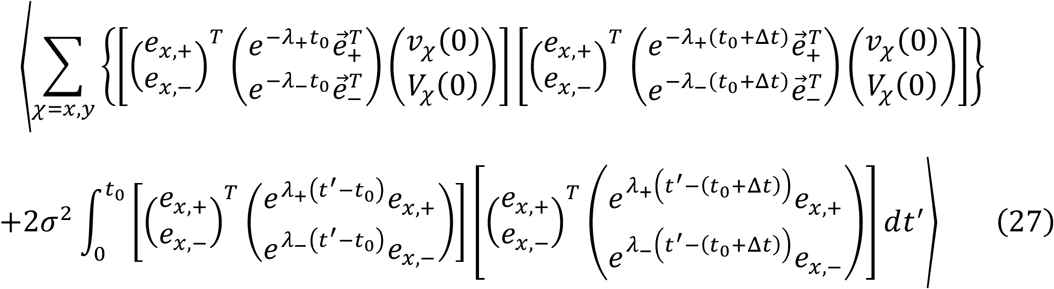

Since *λ*_±_ is always positive when α, β, *γ* > 0, the first term of VAC disappears with time at the rate of 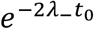. The lower limit of integration also disappears at the same rate. The only time-independent term comes from the upper limit of integration and is the steady state solution of VAC:

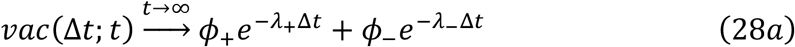

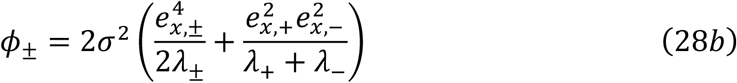

where the second line is another representation of *ϕ*_±_ defined above. Finally, considering the sampling procedure where the velocity is observed as the difference of discretely sampled positions, the representation of *vac*^*ss*^ is obtained by time integration of VAC:

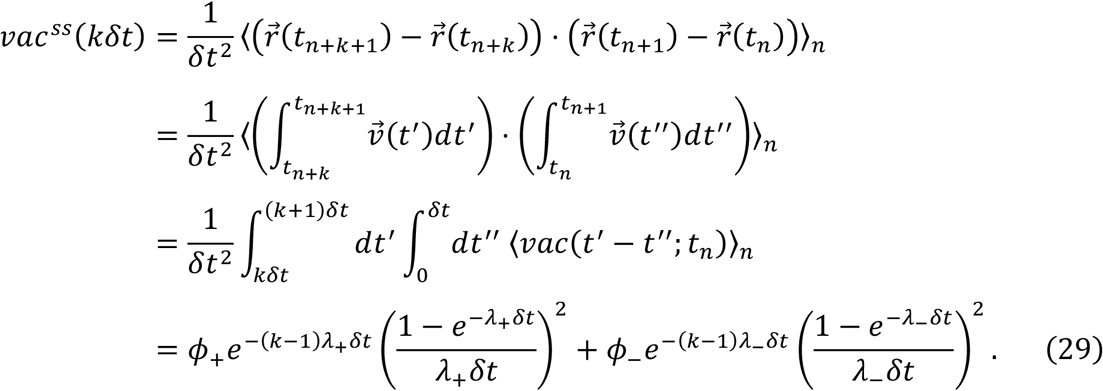

In the third raw, we used the relation 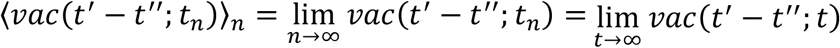 because time average should converge to the steady state solution if the VAC itself converges.

## Author Contributions

SS conceived and designed the work, established the cell culture and contributed to all aspects of data interpretation. MU and YM performed experiments and analyzed the data. AK contributed to the cell mask construction and the centroid analysis. DI performed CNN-based morphology analysis. MU built the automated tracking microscopy and performed majority of the boundary and centroid analysis and model fitting. MU and SS wrote the paper.

## Acknowledgements

The authors thank the present and the past members of the Sawai lab for experimental supports and discussion.

## Supplementary Material

The supplementary Material for this article can be found online at https://www.frontiers.org/articiles/XXXXX

## References

Alirezaeizanjani, Z., Großmann, R., Pfeifer, V., Hintsche, M., and Beta, C. (2020). Chemotaxis strategies of bacteria with multiple run modes. Sci Adv 6, eaaz6153. doi: 10.1126/sciadv.aaz6153.

Ariel, G., Rabani, A., Benisty, S., Partridge, J. D., Harshey, R. M., and Be’er, A. (2015). Swarming bacteria migrate by Lévy Walk. Nat Commun 6, 8396. doi: 10.1038/ncomms9396.

Arzt, M., Deschamps, J., Schmied, C., Pietzsch, T., Schmidt, D., Tomancak, P., et al. (2022). LABKIT: Labeling and Segmentation Toolkit for Big Image Data. Frontiers Comput Sci 4. doi: 10.3389/fcomp.2022.777728.

Bartumeus, F., Catalan, J., Fulco, U. L., Lyra, M. L., and Viswanathan, G. M. (2002). Optimizing the Encounter Rate in Biological Interactions: Lévy versus Brownian Strategies. Phys Rev Lett 88, 097901. doi: 10.1103/physrevlett.88.097901.

Brown, M. W., Silberman, J. D., and Spiegel, F. W. (2012). A contemporary evaluation of the acrasids (Acrasidae, Heterolobosea, Excavata). Eur. J. Protistol. 48, 103–123. doi: 10.1016/j.ejop.2011.10.001.

Chi, Q., Yin, T., Gregersen, H., Deng, X., Fan, Y., Zhao, J., et al. (2014). Rear actomyosin contractility-driven directional cell migration in three-dimensional matrices: a mechano-chemical coupling mechanism. J Roy Soc Interface 11, 20131072 20131072. doi: 10.1098/rsif.2013.1072.

Dieterich, P., Klages, R., Preuss, R., and Schwab, A. (2008). Anomalous dynamics of cell migration. Proc National Acad Sci 105, 459–463. doi: 10.1073/pnas.0707603105.

Driscoll, M. K., McCann, C., Kopace, R., Homan, T., Fourkas, J. T., Parent, C., et al. (2012). Cell shape dynamics: from waves to migration. Plos Comput Biol 8, e1002392. doi: 10.1371/journal.pcbi.1002392.

Dunn, G. A., and Brown, A. F. (1987). A Unified Approach to Analysing Cell Motility. J Cell Sci 1987, 81–102. doi: 10.1242/jcs.1987.supplement_8.5.

Ebata, H., Yamamoto, A., Tsuji, Y., Sasaki, S., Moriyama, K., Kuboki, T., et al. (2018). Persistent random deformation model of cells crawling on a gel surface. Sci Rep-uk 8, 5153. doi: 10.1038/s41598-018-23540-x.

Ershov, D., Phan, M.-S., Pylvänäinen, J. W., Rigaud, S. U., Blanc, L. L., Charles-Orszag, A., et al. (2022). TrackMate 7: integrating state-of-the-art segmentation algorithms into tracking pipelines. Nat. Methods 19, 829–832. doi: 10.1038/s41592-022-01507-1.

Fritz-Laylin, L. K., Lord, S. J., and Mullins, R. D. (2017). WASP and SCAR are evolutionarily conserved in actin-filled pseudopod-based motility. J Cell Biol 216, 1673 1688. doi: 10.1083/jcb.201701074.

Fritz-Laylin, L. K., Prochnik, S. E., Ginger, M. L., Dacks, J. B., Carpenter, M. L., Field, M. C., et al. (2010). The genome of Naegleria gruberi illuminates early eukaryotic versatility. Cell 140, 631–642. doi: 10.1016/j.cell.2010.01.032.

Fujimori, T., Nakajima, A., Shimada, N., and Sawai, S. (2019). Tissue self-organization based on collective cell migration by contact activation of locomotion and chemotaxis. Proc National Acad Sci 116, 4291 4296. doi: 10.1073/pnas.1815063116.

Fulton, C. (1970). Amebo-flagellates as research partners: the laboratory biology of Naegleria and Tetramitus. Methods in Cell Physiology 4, 341–476.

Gail, M. H., and Boone, C. W. (1970). The Locomotion of Mouse Fibroblasts in Tissue Culture. Biophys J 10, 980–993. doi: 10.1016/s0006-3495(70)86347-0.

Haastert, P. J. M. V., and Bosgraaf, L. (2009). Food searching strategy of amoeboid cells by starvation induced run length extension. Plos One 4, e6814. doi: 10.1371/journal.pone.0006814.

Haastert, P. J. M. van (2021). Short- and long-term memory of moving amoeboid cells. Plos One 16, e0246345. doi: 10.1371/journal.pone.0246345.

Harris, T. H., Banigan, E. J., Christian, D. A., Konradt, C., Wojno, E. D. T., Norose, K., et al. (2012). Generalized Lévy walks and the role of chemokines in migration of effector CD8+ T cells. Nature 486, 545 548. doi: 10.1038/nature11098.

Hartman, R., Lau, K., and Chou, W. (1994). The fundamental motor of the human neutrophil is not random: evidence for local non-Markov movement in neutrophils. Biophysical Journal 67.

Huda, S., Weigelin, B., Wolf, K., Tretiakov, K. V., Polev, K., Wilk, G., et al. (2018). Lévy-like movement patterns of metastatic cancer cells revealed in microfabricated systems and implicated in vivo. Nat Commun 9, 4539. doi: 10.1038/s41467-018-06563-w.

Huo, H., He, R., Zhang, R., and Yuan, J. (2021). Swimming Escherichia coli Cells Explore the Environment by Lévy Walk. Appl Environ Microb 87, e02429–20. doi: 10.1128/aem.02429-20.

Imoto, D., Saito, N., Nakajima, A., Honda, G., Ishida, M., Sugita, T., et al. (2021). Comparative mapping of crawling-cell morphodynamics in deep learning-based feature space. Plos Comput Biol 17, e1009237. doi: 10.1371/journal.pcbi.1009237.

Itakura, A., Aslan, J. E., Kusanto, B. T., Phillips, K. G., Porter, J. E., Newton, P. K., et al. (2013). p21-Activated Kinase (PAK) Regulates Cytoskeletal Reorganization and Directional Migration in Human Neutrophils. PLoS ONE 8, e73063. doi: 10.1371/journal.pone.0073063.

Jerison, E. R., and Quake, S. R. (2020). Heterogeneous T cell motility behaviors emerge from a coupling between speed and turning in vivo. Elife 9, e53933. doi: 10.7554/elife.53933.

Kidger, P., Foster, J., Li, X., Oberhauser, H., and Lyons, T. (2021). Neural SDEs as Infinite-Dimensional GANs. Arxiv. doi: 10.48550/arxiv.2102.03657.

King, C. A., Davies, A. H., and Preston, T. M. (1981). Lack of substrate specificity on the speed of amoeboid locomotion inNaegleria gruberi. Experientia 37, 709–710. doi: 10.1007/bf01967936.

King, C. A., Westwood, R., Cooper, L., and Preston, T. M. (1979). Speed of locomotion of the soil amoebaNaegleria gruberi in media of different ionic compositions with special reference to interactions with the substratum. Protoplasma 99, 323–334. doi: 10.1007/bf01275804.

Kuhl, F. P., and Giardina, C. R. (1982). Elliptic Fourier features of a closed contour. Comput Vision Graph 18, 236–258. doi: 10.1016/0146-664x(82)90034-x.

Kwon, T., Kwon, O.-S., Cha, H.-J., and Sung, B. J. (2019). Stochastic and Heterogeneous Cancer Cell Migration: Experiment and Theory. Sci Rep-uk 9, 16297. doi: 10.1038/s41598-019-52480-3.

Li, L., Cox, E. C., and Flyvbjerg, H. (2011). “Dicty dynamics”: Dictyostelium motility as persistent random motion. Phys Biol 8, 046006. doi: 10.1088/1478-3975/8/4/046006.

Li, L., Nørrelykke, S. F., and Cox, E. C. (2008). Persistent Cell Motion in the Absence of External Signals: A Search Strategy for Eukaryotic Cells. Plos One 3, e2093. doi: 10.1371/journal.pone.0002093.

Li, X., Wong, T.-K. L., Chen, R. T. Q., and Duvenaud, D. (2020). Scalable Gradients for Stochastic Differential Equations. Arxiv. doi: 10.48550/arxiv.2001.01328.

Marciano-Cabral, F., and Cline, M. (1987). Chemotaxis by Naegleria fowleri for Bacteria. J. Protozool. 34, 127–131. doi: 10.1111/j.1550-7408.1987.tb03147.x.

Morales, J. C. F., Xue, Q., and Roh-Johnson, M. (2022). An evolutionary and physiological perspective on cell-substrate adhesion machinery for cell migration. Frontiers Cell Dev Biology 10, 943606. doi: 10.3389/fcell.2022.943606.

Nakajima, A., Ishida, M., Fujimori, T., Wakamoto, Y., and Sawai, S. (2016). The microfluidic lighthouse: an omnidirectional gradient generator. Lab Chip 16, 4382–4394. doi: 10.1039/c6lc00898d.

Nakamura, K., and Kobayashi, T. J. (2021). Connection between the Bacterial Chemotactic Network and Optimal Filtering. Phys Rev Lett 126, 128102. doi: 10.1103/physrevlett.126.128102.

Ohta, T., Tarama, M., and Sano, M. (2016). Simple model of cell crawling. Phys D Nonlinear Phenom 318–319, 3–11. doi: 10.1016/j.physd.2015.10.007.

Parfrey, L. W., Lahr, D. J. G., Knoll, A. H., and Katz, L. A. (2011). Estimating the timing of early eukaryotic diversification with multigene molecular clocks. Proc National Acad Sci 108, 13624–13629. doi: 10.1073/pnas.1110633108.

Partridge, B., Anton, S. G., Khorshed, R., Adams, G., Pospori, C., Celso, C. L., et al. (2022). Heterogeneous run-and-tumble motion accounts for transient non-Gaussian super-diffusion in haematopoietic multi-potent progenitor cells. Plos One 17, e0272587. doi: 10.1371/journal.pone.0272587.

Passucci, G., Brasch, M. E., Henderson, J. H., Zaburdaev, V., and Manning, M. L. (2019). Identifying the mechanism for superdiffusivity in mouse fibroblast motility. Plos Comput Biol 15, e1006732. doi: 10.1371/journal.pcbi.1006732.

Pollard, T. D. (2007). Regulation of actin filament assembly by Arp2/3 complex and formins. Annu Rev Bioph Biom 36, 451 477. doi: 10.1146/annurev.biophys.35.040405.101936.

Preston, T. M., and King, C. A. (1978). An experimental study of the interaction between the soil amoeba Naegleria gruberi and a glass substrate during amoeboid locomotion. J Cell Sci 34, 145–158. doi: 10.1242/jcs.34.1.145.

Prostak, S. M., Robinson, K. A., Titus, M. A., and Fritz-Laylin, L. K. (2021). The actin networks of chytrid fungi reveal evolutionary loss of cytoskeletal complexity in the fungal kingdom. Curr Biol 31, 1192–1205.e6. doi: 10.1016/j.cub.2021.01.001.

Reynolds, A. M. (2010). Can spontaneous cell movements be modelled as Lévy walks? Phys Statistical Mech Appl 389, 273–277. doi: 10.1016/j.physa.2009.09.027.

Reynolds, A. M., and Ouellette, N. T. (2016). Swarm dynamics may give rise to Lévy flights. Sci Rep-uk 6, 30515. doi: 10.1038/srep30515.

Rieu, J.-P., Delanoë-Ayari, H., Takagi, S., Tanaka, Y., and Nakagaki, T. (2015). Periodic traction in migrating large amoeba of Physarum polycephalum. J Roy Soc Interface 12, 20150099 20150099. doi: 10.1098/rsif.2015.0099.

Rodiek, B., and Hauser, M. J. B. (2015). Migratory behaviour of Physarum polycephalum microplasmodia. Eur. Phys. J. Spéc. Top. 224, 1199–1214. doi: 10.1140/epjst/e2015-02455-2.

Sadhu, R. K., Iglič, A., and Gov, N. S. (2023). A minimal cell model for lamellipodia-based cellular dynamics and migration. J. Cell Sci. 136. doi: 10.1242/jcs.260744.

Sebé-Pedrós, A., Grau-Bové, X., Richards, T. A., and Ruiz-Trillo, I. (2014). Evolution and Classification of Myosins, a Paneukaryotic Whole-Genome Approach. Genome Biol. Evol. 6, 290–305. doi: 10.1093/gbe/evu013.

Selmeczi, D., Li, L., Pedersen, L. I. I., Nrrelykke, S. F., Hagedorn, P. H., Mosler, S., et al. (2008). Cell motility as random motion: A review. European Phys J Special Top 157, 1–15. doi: 10.1140/epjst/e2008-00626-x.

Selmeczi, D., Mosler, S., Hagedorn, P. H., Larsen, N. B., and Flyvbjerg, H. (2005). Cell motility as persistent random motion: theories from experiments. Biophys J 89, 912 931. doi: 10.1529/biophysj.105.061150.

Shaebani, M. R., Jose, R., Santen, L., Stankevicins, L., and Lautenschläger, F. (2020). Persistence-Speed Coupling Enhances the Search Efficiency of Migrating Immune Cells. Phys Rev Lett 125, 268102. doi: 10.1103/physrevlett.125.268102.

Shaebani, M. R., Piel, M., and Lautenschläger, F. (2022). Distinct speed and direction memories of migrating dendritic cells diversify their search strategies. Biophys J 121, 4099–4108. doi: 10.1016/j.bpj.2022.09.033.

Stokes, C. L., Lauffenburger, D. A., and Williams, S. K. (1991). Migration of individual microvessel endothelial cells: stochastic model and parameter measurement. J Cell Sci 99, 419–430. doi: 10.1242/jcs.99.2.419.

Takagi, H., Sato, M. J., Yanagida, T., and Ueda, M. (2008). Functional Analysis of Spontaneous Cell Movement under Different Physiological Conditions. Plos One 3, e2648. doi: 10.1371/journal.pone.0002648.

Taktikos, J., Stark, H., and Zaburdaev, V. (2013). How the Motility Pattern of Bacteria Affects Their Dispersal and Chemotaxis. Plos One 8, e81936. doi: 10.1371/journal.pone.0081936.

Thong, Y. H., and Ferrante, A. (1986). Migration patterns of pathogenic and nonpathogenic Naegleria spp. Infection and Immunity 51, 177 180.

Tweedy, L., Meier, B., Stephan, J., Heinrich, D., and Endres, R. G. (2013). Distinct cell shapes determine accurate chemotaxis. Sci Rep-uk 3, 2606. doi: 10.1038/srep02606.

Velle, K. B., and Fritz-Laylin, L. K. (2019). Diversity and evolution of actin-dependent phenotypes. Curr. Opin. Genet. Dev. 58, 40–48. doi: 10.1016/j.gde.2019.07.016.

Velle, K. B., and Fritz-Laylin, L. K. (2020). Conserved actin machinery drives microtubule-independent motility and phagocytosis in Naegleria. J Cell Biology 219, e202007158. doi: 10.1083/jcb.202007158.

Viswanathan, G. M., Buldyrev, S. V., Havlin, S., Luz, M. G. E. da Raposo, E. P., and Stanley, H. E. (1999). Optimizing the success of random searches. Nature 401, 911–914. doi: 10.1038/44831.

Walsh, C. J. (2007). The role of actin, actomyosin and microtubules in defining cell shape during the differentiation of Naegleria amebae into flagellates. Eur J Cell Biol 86, 85–98. doi: 10.1016/j.ejcb.2006.10.003.

Welf, E. S., Ahmed, S., Johnson, H. E., Melvin, A. T., and Haugh, J. M. (2012). Migrating fibroblasts reorient directionality by a metastable, PI3K-dependent mechanism. J Cell Biology 197, 105 114. doi: 10.1083/jcb.201108152.

Xu, X., Wen, X., Veltman, D. M., Keizer-Gunnink, I., Pots, H., Kortholt, A., et al. (2017). GPCR-controlled membrane recruitment of negative regulator C2GAP1 locally inhibits Ras signaling for adaptation and long-range chemotaxis. Proc National Acad Sci 114, E10092–E10101. doi: 10.1073/pnas.1703208114.

