## Supplementary Materials for "Random walk and cell morphology dynamics in *Naegleria gruberi*"

*** Correspondence:**

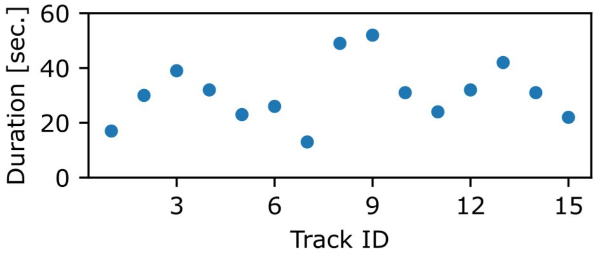


**Supplementary Figure S1 |** Duration of membrane protrusions. Protrusions were manually tracked from its formation to disappearance. The mean duration was 31 seconds.

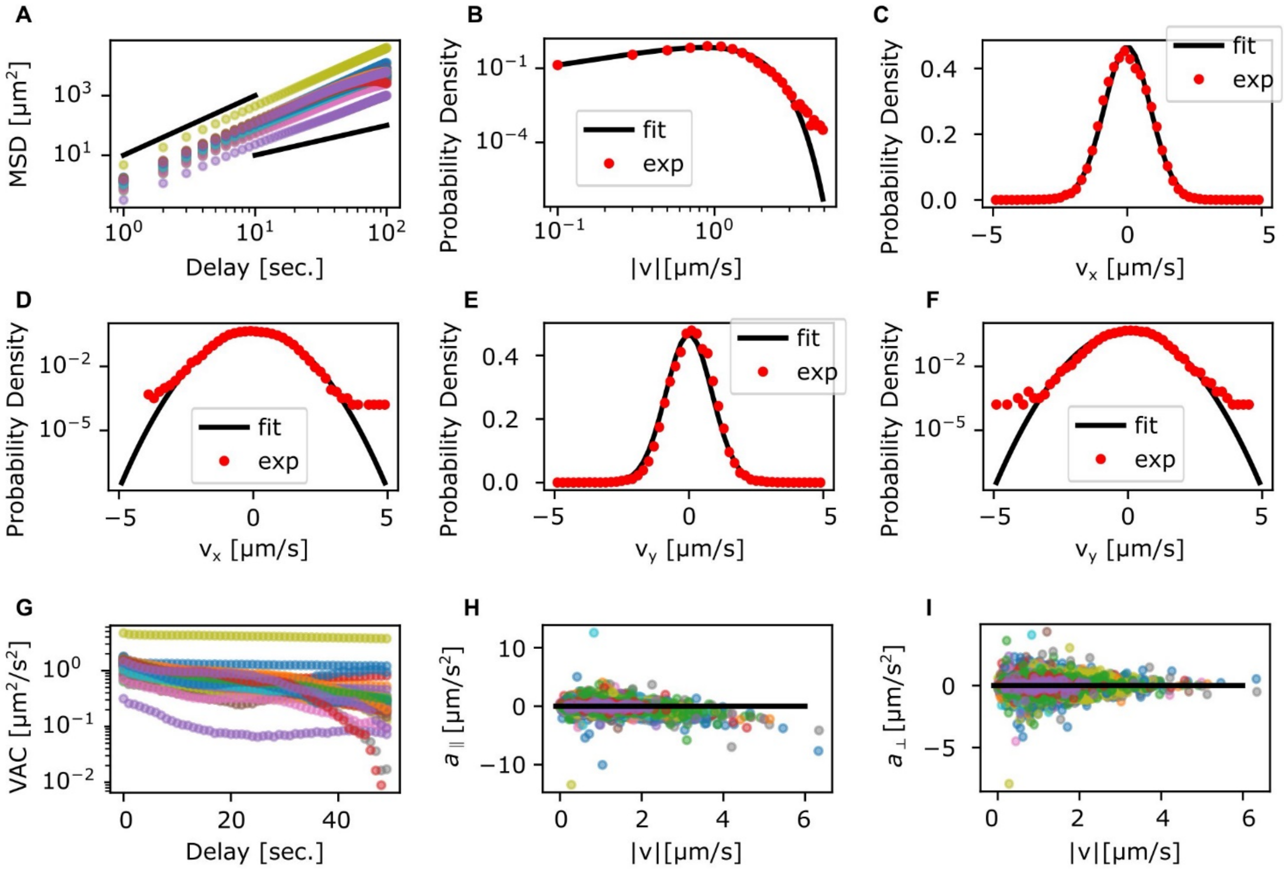


**Supplementary Figure S2 |** Detailed statistics of *N. gruberi* cell migration. (**A**) MSD of individual trajectories. (**B-F**) Probability distribution of velocity. Velocity ( $|\hat{\vec{v}}|$ ) distribution in log-log scale (B). $\hat{v}_x$ (C, D) and $\hat{v}_y$ (E, F) in linear (C, E) or semi-log (D, F) scale. (**G**) VAC of individual trajectories. (**H**, **I**) Scatter plot of acceleration versus absolute speed. $a_{\perp}$ (H) and $a_{\parallel}$ (I). Colors represent each trajectory.

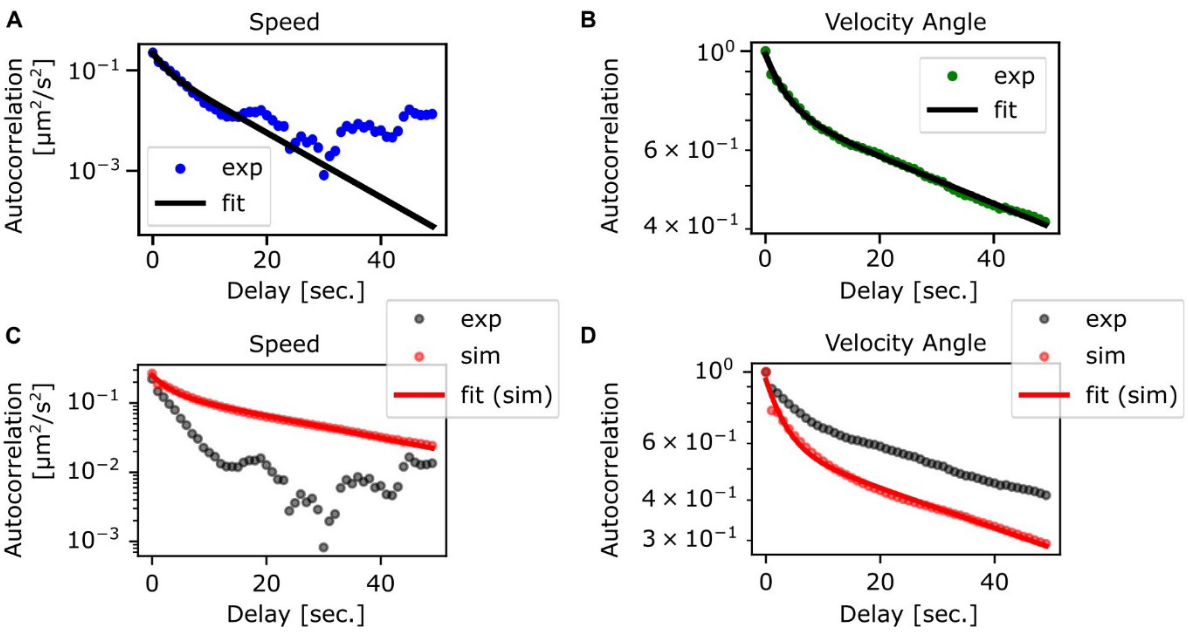


**Supplementary Figure S3 |** Detailed analysis of velocity autocorrelation. (**A**) Autocorrelation of the absolute speed $|\hat{\vec{v}}|$. Blue circles: experiments. Black curve: fitting curve with double exponential function. (**B**) Autocorrelation of the velocity angle. Green circles: autocorrelation of $\hat{\vec{v}}/|\hat{\vec{v}}|$. Black curve: fitting curve with double exponential function. (**C**) Autocorrelation of the absolute speed from the generalized Langevin equation with positional uncertainty. Red circle: simulation. Red curve: fitting curve with double exponential. Gray circle: experiment (see **Supplementary Table S1A**, **B**). (**D**) Autocorrelation of the velocity angle. Red circle: $\hat{\vec{v}}/|\hat{\vec{v}}|$ from simulation. Red curve: the fitting curve with double exponential function. Gray circle: experiment (see also **Supplementary Table S1C, D**).

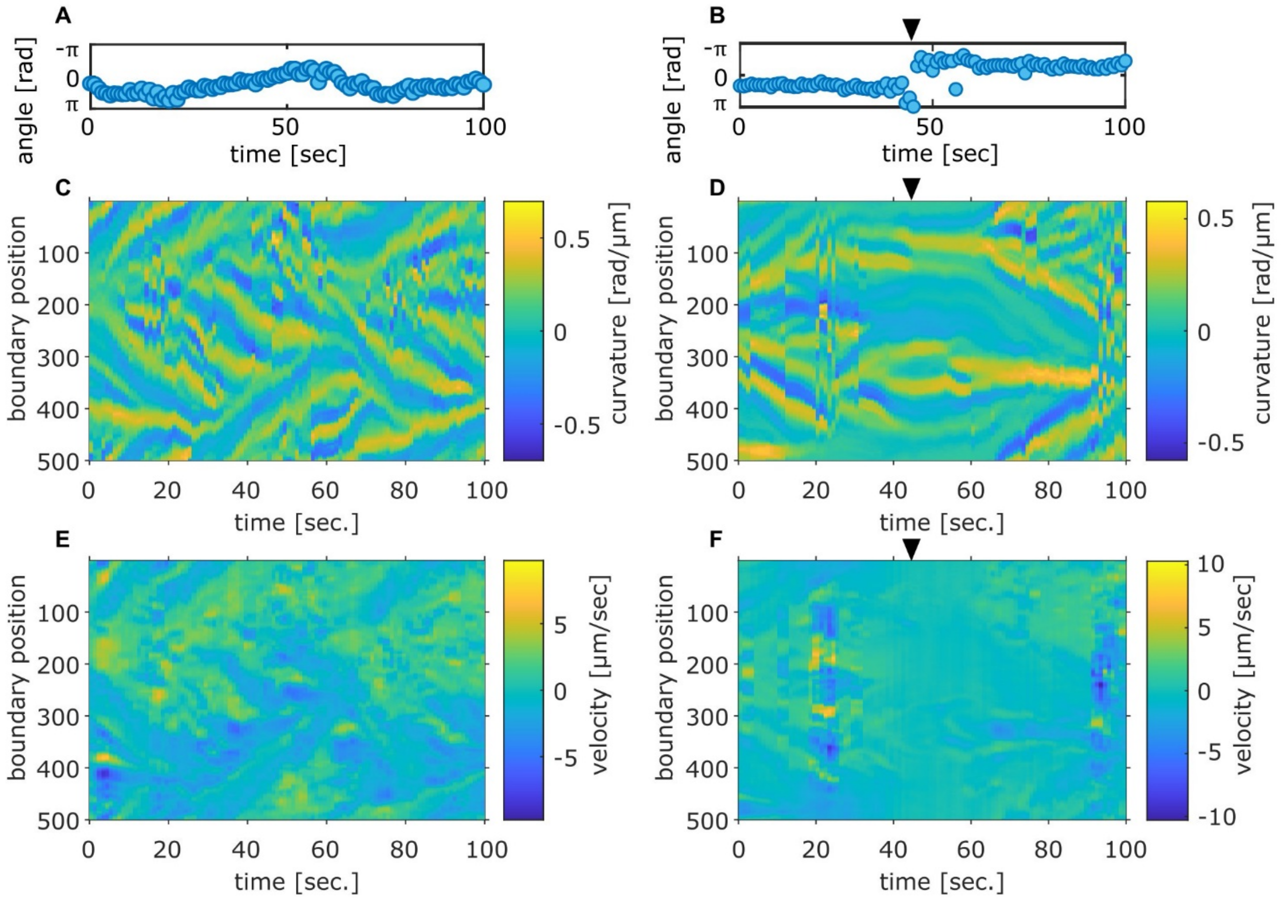


**Supplementary Figure S4 |** Comparison between the centroid velocity angle and the boundary curvature and velocity. Panels (A, C, E) are from a representative time series. Panels (B, D, F) are from a time series which showed dumbbell shape and turning of centroid velocity at the time indicated by black arrows. (**A**, **B**) Time evolution of the centroid velocity angle. (**C**, **D**) Boundary curvature. (**E**, **F**) Kymographs of the velocity on the cell mask edge.

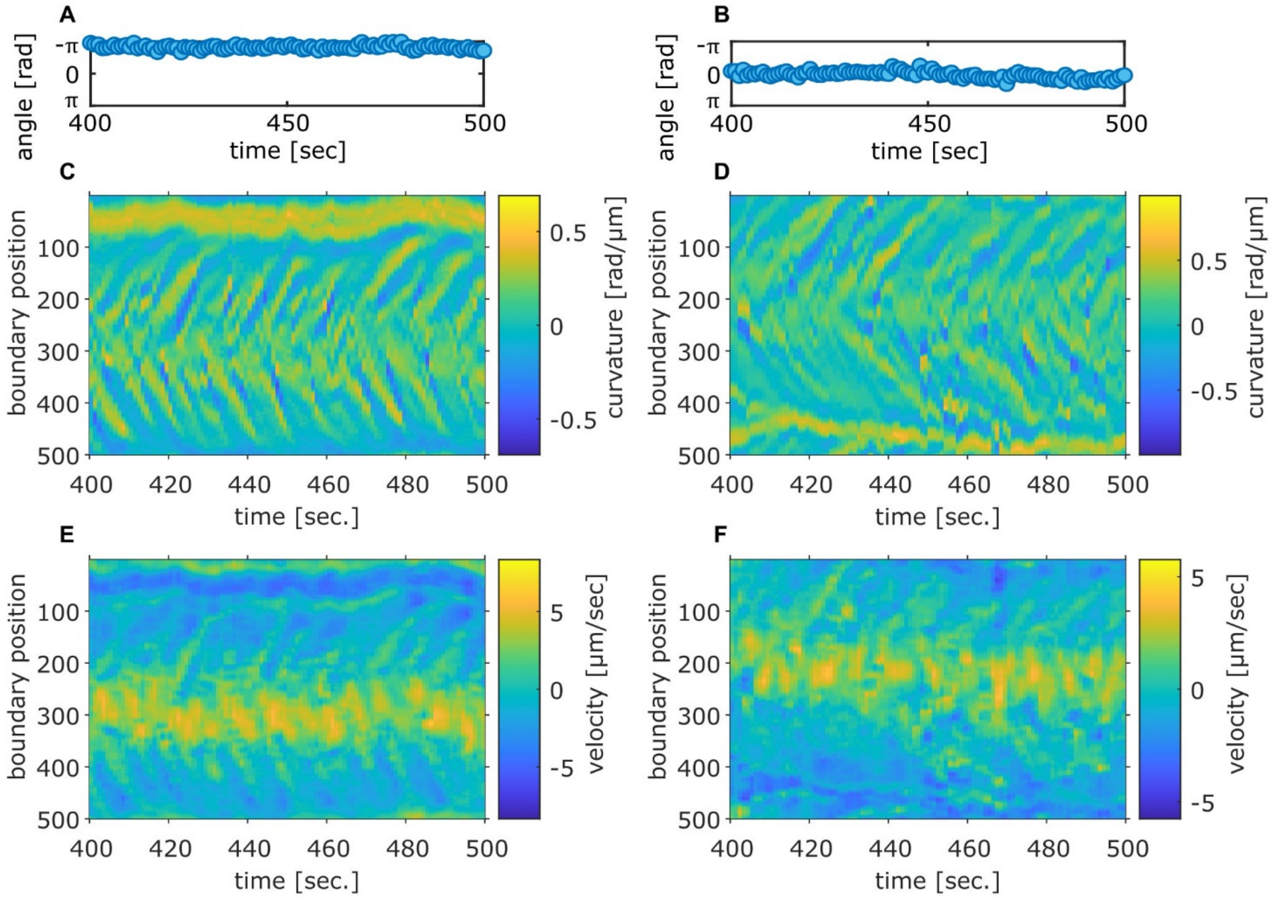


**Supplementary Figure S5 |** The centroid velocity angle, the curvature and local velocity in cells exhibiting persistent centroid movement. Two independent samples (A, C, E) and (B, D, F). (**A**, **B**) Time evolution of the centroid velocity angle. (**C**, **D**) The edge curvature. (**E**, **F**) The normal velocity along the cell boundary.

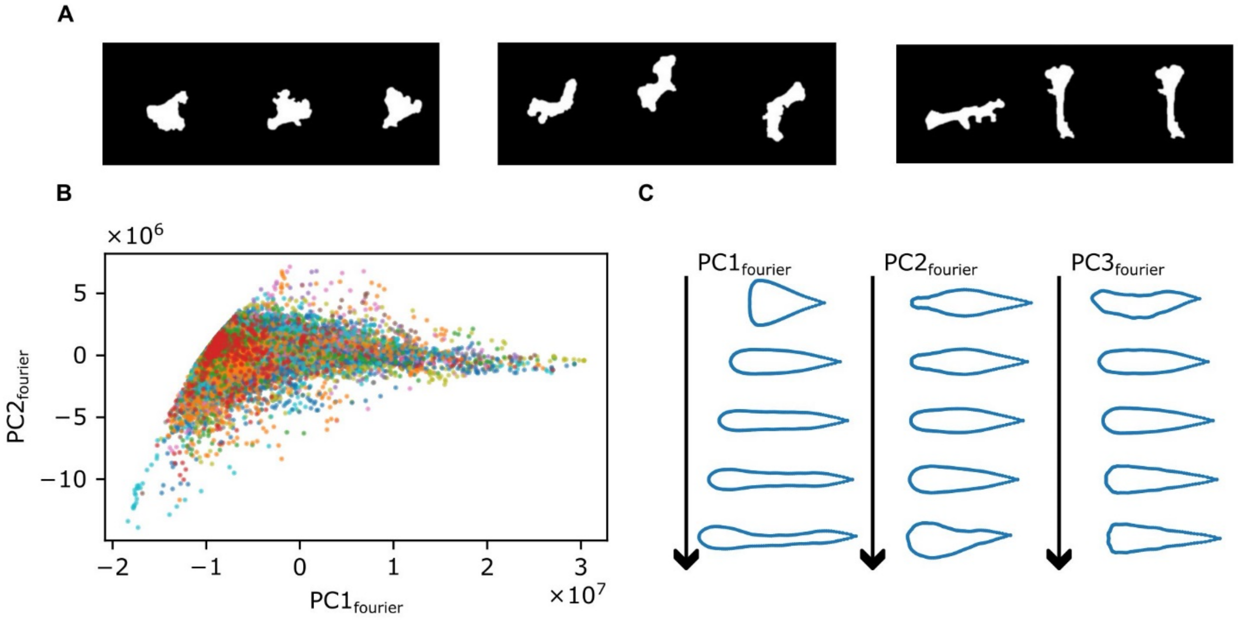


**Supplementary Figure S6 | Fourier-descriptor based cell shape analysis**. (**A**) Representative masks chosen by eye. Three representative masks of fan-shape (left), split (center), stick-like shape (right). (**B**) Scatter plot of all masks. Different time series are indicated by different color. (**C**) Reverse-transformed shape from sets of arbitrary principal component values. $PC1_{fourier}$ (top), $PC2_{fourier}$ (middle), or $PC3_{fourier}$ (bottom) values were varied. Other principal components were kept at 0.

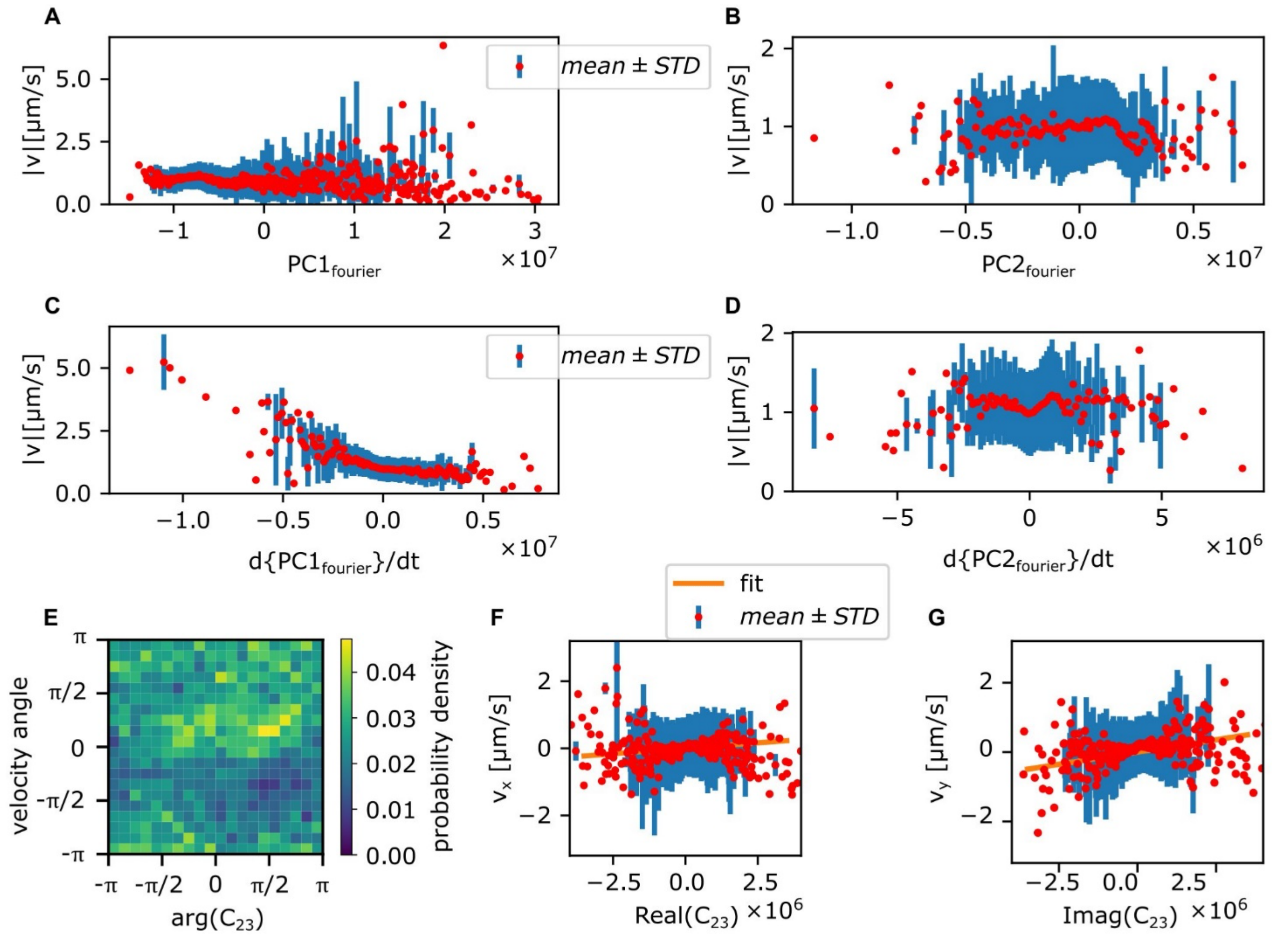


**Supplementary Figure S7 |** Comparison of cell shape and centroid velocity. (**A, B**) Centroid speed is plotted as a function of $PC1_{fourier}$ (A) and $PC2_{fourier}$ (B). (**C, D**) Centroid speed is plotted as a function of $d\{PC1_{fourier}\}/dt$ (C) and $d\{PC2_{fourier}\}/dt$ (D). (**E, F, G**) Cell shape dynamics in elongation and triangle modes is compared with centroid velocity. (**E**) Distribution of angles of centroid velocity and $C_{23}$. (**F**, **G**) The x- (**F**) and y-components (**G**) of centroid velocity plotted against real (**F**) and imaginary (**G**) parts of $C_{23}$. Red circles and blue bars indicate average and standard deviation of centroid velocity binned with the value of $C_{23}$. Orange lines indicate the result of fitting with linear proportionality.

**Supplementary Table S1.** Parameter values obtained by fitting the autocorrelations in Supplementary Figure S3 with double exponential. The autocorrelations of (A, B) the absolute speed from experiment (A) or simulation (B), (C,D) the velocity angle from experiment (C) or simulation (D) were fit. These autocorrelations were fit with $Autocorrelation = \Phi_1^{sp} e^{-\delta t/T_1^{sp}} + \Phi_2^{sp} e^{-\delta t/T_2^{sp}}$. Each row in the table shows the obtained parameter set. The unit of $\Phi_1^{sp}$ and $\Phi_2^{sp}$ are [$\mu m^2/s^2$] for rows (A,B) and [A.U.] for rows (C,D).

| | $\boldsymbol{T_1^{sp}}$ $[\boldsymbol{sec.}]$ | $\boldsymbol{T_2^{sp}}$ $[\boldsymbol{sec.}]$ | $\boldsymbol{\Phi_1^{sp}}$ | $\boldsymbol{\Phi_2^{sp}}$ |
|---|---|---|---|---|
| (A) | 1.8 | 6.8 | 0.11 | 0.11 |
| (B) | 2.7 | 27.0 | 0.12 | 0.14 |
| (C) | 3.7 | 82.5 | 0.24 | 0.74 |
| (D) | 3.3 | 70.4 | 0.37 | 0.58 |

**Supplementary Movie S1**

The movie corresponds to **Figure 1A**. By repositioning the timelapse images (gray) according to the trajectory of automated stage, the movie in the laboratory frame was obtained.

**Supplementary Movie S2**

The dynamics of protrusions. White circles indicate the protrusions tracked from appearance to disappearance to obtain the distribution of lifetime shown in **Supplementary Figure S1**.

**Supplementary Movie S3**

The movie corresponds to **Figure 2A**.

**Supplementary Movie S4**

The movie corresponds to **Figure 2B**.

**Supplementary Movie S5**

The movie corresponds to **Figure 2C**.

**Supplementary Movie S6**

The movie corresponds to **Figure 5C**.
